# Disease Modifying Osteoarthritis Drug Discovery Using A Temporal Phenotypic Reporter In 3D Aggregates of Primary Human Chondrocytes

**DOI:** 10.1101/2023.03.10.532073

**Authors:** Maria A. Cruz, Scott Gronowicz, Makan Karimzadeh, Kari Martyniak, Ramapaada Medam, Thomas J. Kean

## Abstract

Osteoarthritis is a significant and growing problem with no disease modifying drugs in the clinic. Current screening platforms typically use 2D culture, immortalized or non-human cells in a hyperoxic environment. To challenge this paradigm and identify new drugs, we engineered primary human chondrocytes with a secreted luciferase reporter under the control of the articular cartilage marker, type II collagen. We then successfully screened a natural product library using a high throughput model with COL2A1-Gaussia luciferase primary human chondrocyte reporter cells in 3D aggregates under physioxia. We identified several candidate compounds that increased type II collagen over controls, with aromoline being the best candidate. Aromoline is a bisbenzylisoquinoline alkaloid that has been studied for its anti-proliferative, anti-inflammatory, and anti-microbial properties, and we are the first to explore its effects on chondrocytes and chondrogenesis. In silico analysis of predicted targets narrowed by RNA-Seq data on expression provided an unexpected initial candidate target protein: the dopamine receptor D4 (DRD4). The researchers confirmed upregulation in the expression of DRD4 and type II collagen after treatment with aromoline. This novel approach combining in silico and in vitro methods provides a platform for drug discovery in a challenging and under-researched area. In conclusion, a novel drug (aromoline) and target receptor (dopamine receptor D4) were identified as stimulating type II collagen, with the goal to treat or prevent osteoarthritis.

## INTRODUCTION

Osteoarthritis (OA) is the most prevalent articular joint disease worldwide. It affects approximately 16% of adult in the US, with approximately 32.5 million adults reporting OA between 2008 to 2014 ^1^. Post-traumatic osteoarthritis (PTOA) is thought to account for up to 12% of cases ^2^. Current pharmaceutical therapies for OA are primarily palliative including non-steroidal anti-inflammatories, opioid analgesics, intra-articular corticosteroids and hyaluronan injections ^3^. Surgical intervention is the only current treatment that can restore at least partial function to the joint, however results are highly variable ^4^. There are currently no disease-modifying drugs available to treat OA, as such there is a dire need to identify novel regenerative pharmaceutical alternatives ^3^.

Cartilage degeneration is the primary pathological feature of osteoarthritis. Cartilage is composed of a dense extracellular matrix interspersed with chondrocytes. Chondrocytes synthesize a combination of glycosaminoglycans (GAG), proteoglycans and collagens that make up native cartilage. Type II collagen makes up 90-95% of all collagens found in articular cartilage, and its degradation is one of the early symptoms of OA ^5–7^. Furthermore advances in tissue engineering for the treatment of OA and post traumatic osteoarthritis (PTOA) have been hampered by the inability to recapitulate native levels of type II collagen *in vitro* ^8^. Previous work by our group has shown that increased type II collagen synthesis correlates with improved mechanical and biochemical properties in engineered tissue *in vitro* ^9–11^. In this study we propose using type II collagen expression as a chondrogenic phenotype for drug discovery.

Traditional methods to measure type II collagen, such as biochemical assays, protein assays, and real time qPCR are laborious, costly, time consuming, and destructive to samples ^12^. We have developed a phenotypic COL2A1-*Gaussia* luciferase primary human chondrocyte reporter (HuCol2gLuc) that allows for non-destructive temporal analysis of type II collagen production ^13^. In this work, the modified chondrocytes are used in a 3D aggregate model enabling high throughput analysis. Published findings by our group have shown the success of this assay, by using it to identify vitamins and minerals that improve chondrogenesis in the murine chondrogenic cell line, ATDC5, and in primary rabbit chondrocytes ^10, 14^. This system has also been used to identify biomaterials that improve type II collagen production in human primary chondrocytes in 3D bioprinted constructs ^13^. In this study we used the phenotypic reporter system to identify new lead compounds, that can stimulate type II collagen production in primary human chondrocytes by screening the NCI natural products Set V library ^15^.

Natural Products are great source of potential new drugs and lead compounds. This class of compounds have large structural diversity and are enriched in bioactive compounds covering a wider chemical space compared to chemically synthesized compounds ^16, 17^. They have been a prolific source of leads in drug discovery, with several marketed drugs being or having been modified from natural products ^18, 19^. Despite their advantages, isolation and characterization of natural products from their original source, followed by lack of high throughput methodologies to screen natural products, are primary drawbacks that have hampered natural product advancement in drug discovery ^20, 21^.

Using our 3D *in vitro* phenotypic reporter assay we have successfully screened the Natural Product Set V library from NCI for compounds with a statistically significant stimulatory effect on type II collagen production compared to controls. This library consists of 390 compounds selected from the 140,000 products from the DTP Open Repository collection. Compounds for this library were selected based on origin, purity (>90% by ELSD, major peak has the correct mass ion), structural diversity and availability of compound. From the screen we have identified a hit for increased type II collagen production, representing a potential target for OA drug discovery.

## RESULTS

### Initial screen of natural products library identified two hits that increased type II collagen synthesis in vitro

To screen 390 natural products from the NCI natural products library, primary human chondrocytes were modified to express a *Gaussia* luciferase secreted reporter driven by type II collagen promoter. HuCol2gLuc primary human chondrocytes have been previously characterized by our group as retaining chondrogenic capacity and luminescence serving as a proxy for type II collagen expression ^13^. Aggregate cultures of HuCol2gLuc cells were seeded in the presence of the 390 compounds from the natural product library at 5µM continuously over 22 days in defined chondrogenic differentiation medium with feedings every 2-3 days ^22, 23^. **Figure 1A** shows overall results of treatment. Of the 390 compounds, 77 were cytotoxic to cells. These were identified by a failure to form aggregates after initial treatment. Of those that were not cytotoxic 211 compounds were detrimental to chondrogenesis, classified as less than 90% of vehicle (DMSO) control aggregate luminescence on day 22 (**Fig. 1A**). We identified 70 compounds that had little effect on type II collagen expression (90-110% of luminescence) and 32 compounds that were considered beneficial (>110% of control). Of those 32, only two candidates, identified as 84 (aromoline) and 204 (deserpidine), had a statistically significant increase in luminescence as compared to the controls by day 22, shown by the volcano plot in **Fig 1B**. Luminescence over 22 days normalized to our DMSO control is shown in **Fig 1C**. The overall trend of expression for both candidates differs over the 22 days. As seen in **Fig 1C** aggregate sample treated with aromoline shows an increased luminescence over the control as early as day 3 with a sustained increase in expression over 22 days. Alternatively, luminescence of aggregate treated with deserpidine shows an initial decrease in expression with continued increase in luminescence after day 8. Immunohistochemistry of sections collected from aggregates at day 22 shown in **Fig 1D** confirm expression of type II collagen in HuCol2gLuc derived aggregates with substantially darker staining noted in the aggregate treated with aromoline.

**Fig. 1.**
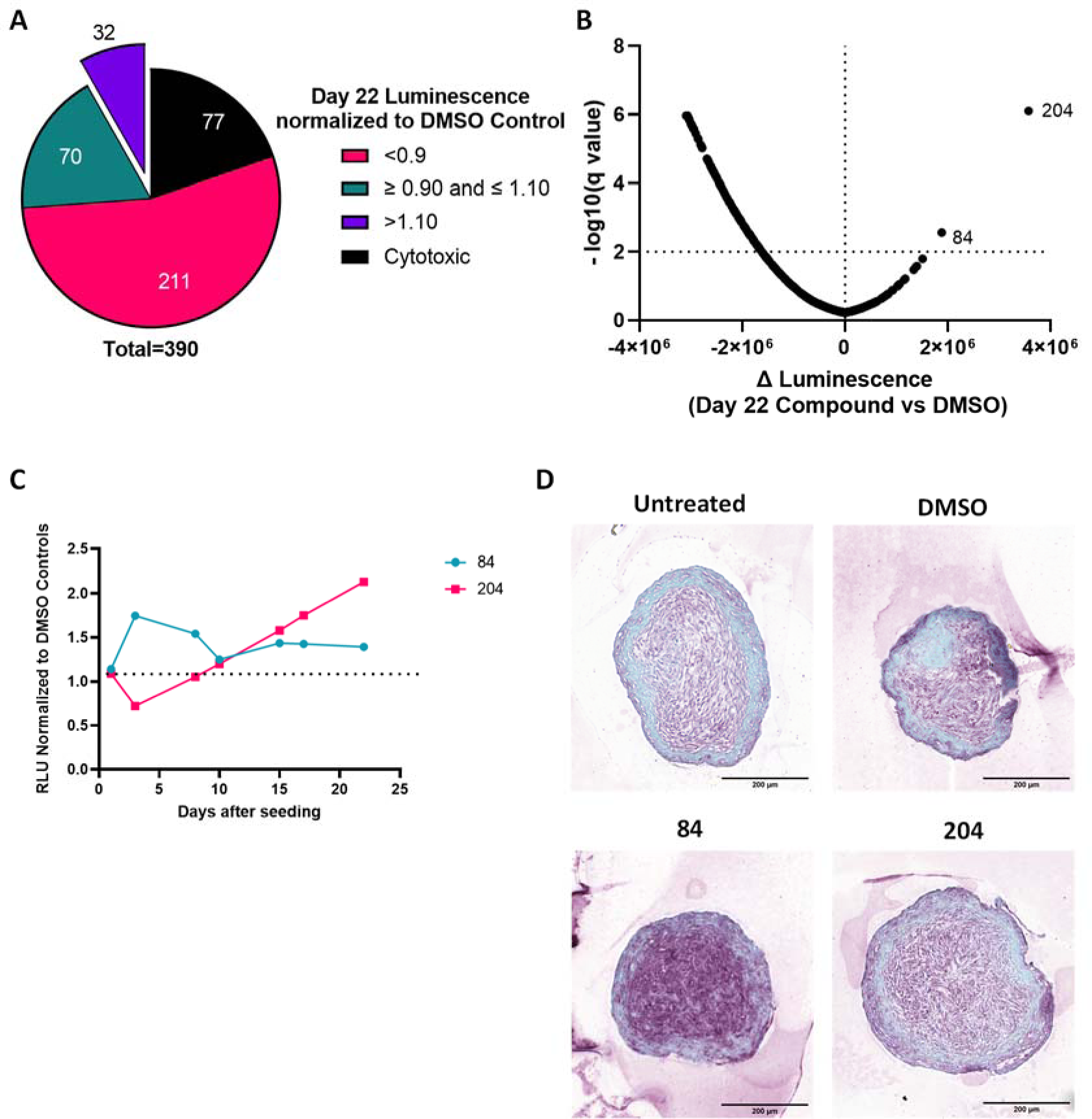
Drug Screen of natural product library via HuCol2gLuc system identified hits that increased type II collagen. (**A**) Overview of Day 22 luminescence for 390 Natural Drug Screen Library normalized to DMSO (vehicle) control. (**B**) Volcano Plot of day 22 aggregate luminescence vs. DMSO control. (**C**) Temporal luminescence signal of aggregates treated with candidates 84 (aromoline) and 204 (deserpidine) over 22 days with results normalized to DMSO control. (**D**) Immunohistological staining for type II collagen of day 22 aggregates treated with candidate 84, 204 and controls for type II collagen. Scale Bars = 200 µm.

### Validation of hits from the screen in three different cell donors verified aromoline as a hit

To validate aromoline and deserpidine from the drug screen, HuCol2gLuc cells, from three different donors, were treated with either aromoline or deserpidine in aggregate culture for 22 days in chondrogenic differentiation medium supplemented with 10ng/ml of TGF-β1. As seen in **Fig 2A, 2C** and **2E**, deserpidine treatment resulted in an increase in luminescence compared to the untreated group in cell aggregates from donors 1 and 3, but not donor 2. Luminescence for deserpidine in donor 1 aggregates decreased from day 0 to day 10 with a significant increase from day 15 to day 22, similar to what was observed in the initial drug screen while donor 3 had a significant increase on days 9 and 14 with little difference early or late in the experiment. In contrast, aromoline resulted in increased luminescence for donors 2 and 3 both cumulatively and on days 8 and 10 for donor 1 and days 5,7,9 and 14 for donor 3, shown in **Fig 2C-2F**, but only at day 3 for donor 1, shown in **Fig 2A**. The increase in luminescence for donor 2 aggregates treated with aromoline compared to the untreated control occurred from day 0 to day 10 with no change from day 15 to day 22, as compared to the control (**Fig 2C)**.

**Fig. 2.**
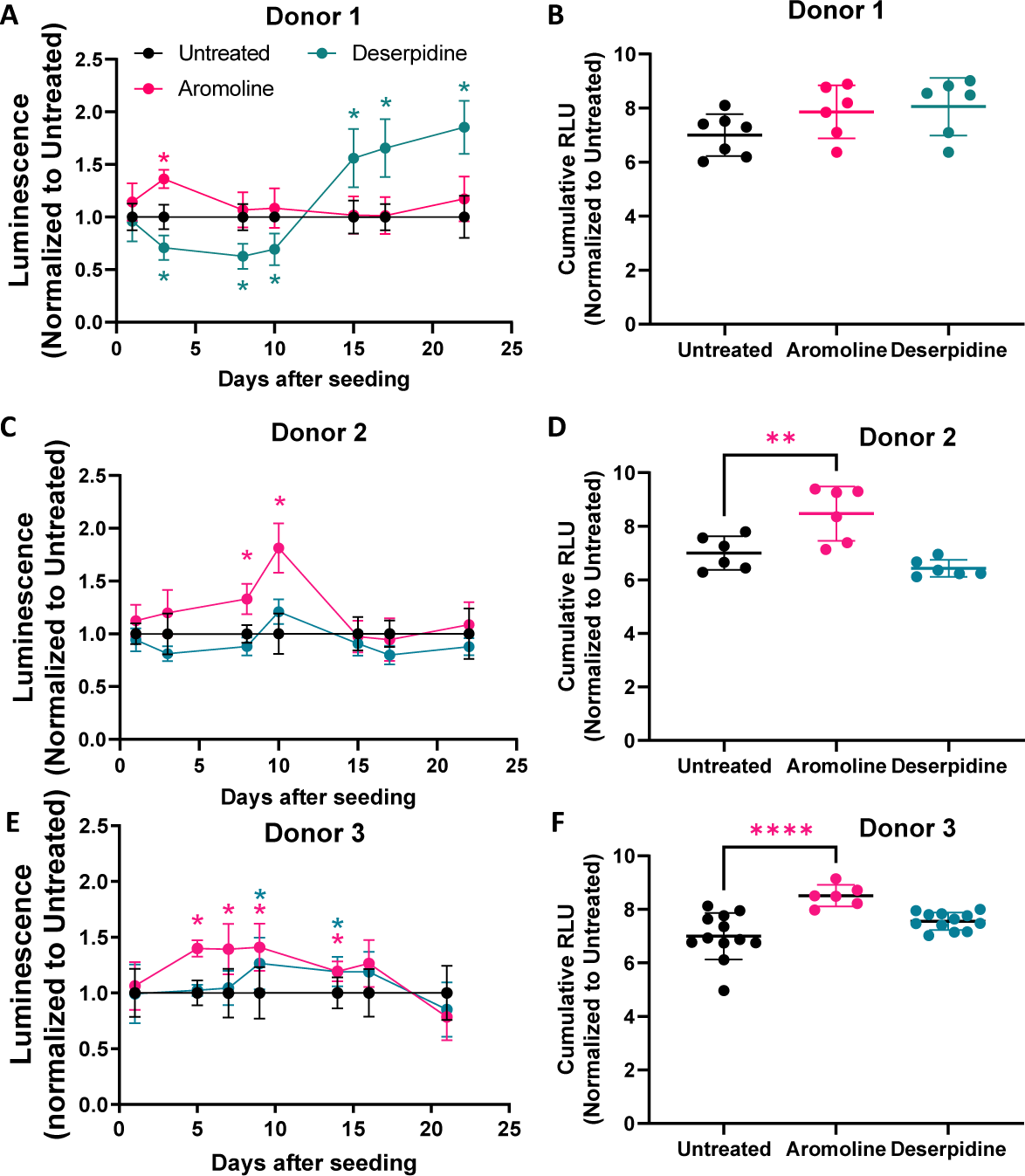
Validation of hits confirmed increased type II collagen expression with aromoline treatment. Drug Candidates selected from the screen were tested in aggregate culture of HuCol2gLuc reporter human chondrocytes from three different donors. Results are shown as luminescence over 22 days normalized to an untreated control, for donors 1,2 and 3 (**A,C,E**). Results are also presented as cumulative luminescence signal for donors 1,2 and 3 (**B,D,F**). N ≥ 6. Individual replicates or mean of replicates are shown with error bars indicating standard deviation and * indicating p <0.01 vs. untreated control, * aromoline; * deserpidine.

This is also similar to the initial screen, where aromoline had a peak in luminescence early in chondrogenesis versus late. Contrary to **Fig 2A**, deserpidine showed no significant increase in donor 1 for cumulative luminescence. As seen in **Fig 2D**, aromoline showed a significant increase in cumulative luminescence for donor 2 with a positive trend for donor 1.

To corroborate luminescence results, biochemical assays to quantitate DNA, glycosaminoglycan (GAG) and total collagen content were performed at the end of culture (day 22). **Fig. 3A and 3D** shows an average total DNA content of ∼0.2 μg of per donor 1 samples and ∼0.7 μg per donor 2 samples. There is no significant difference between drug treatments for donor 1. Treatment with aromoline also had no change in DNA content for donor 2, however treatment with deserpidine showed a significant decrease in DNA content compared to the untreated control (**Fig. 3D**). This suggests that aromoline has no effect on cell proliferation or viability over 22 days, while deserpidine seems to have a donor dependent effect on cell viability. Total glycosaminoglycan content is shown in **Fig. 3B and 3E**. There seems to be a decrease in GAG content with aromoline in donor 1 and 2 aggregates, although donor 1 results were not statistically significant. Deserpidine treatment had no effect on GAG for donor 1 but had a significant decrease in donor 2 to 10 μg vs. 15 μg in the untreated controls, corroborating the donor dependent effects of deserpidine. Quantification of total collagen, as seen in **Fig. 3C and F** shows no significant increase in total collagen with deserpidine treatment in either donor 1 or donor 2 aggregates vs. untreated control. Treatment with aromoline resulted in a significant increase in donor 2 aggregate collagen, to ∼21 μg over untreated (∼14μg), and a positive upward trend in donor 1. Furthermore, a substantial difference in the total DNA, GAG, and collagen content was noted between donor 1 and donor 2 regardless of treatment. Altogether aggregates from donor 1 cells seem to be less chondrogenic than aggregates from donor 2. Based on cumulative luminescence in donor 2 and total collagen content, aromoline was identified as a top hit for the stimulation of type II collagen.

**Fig. 3.**
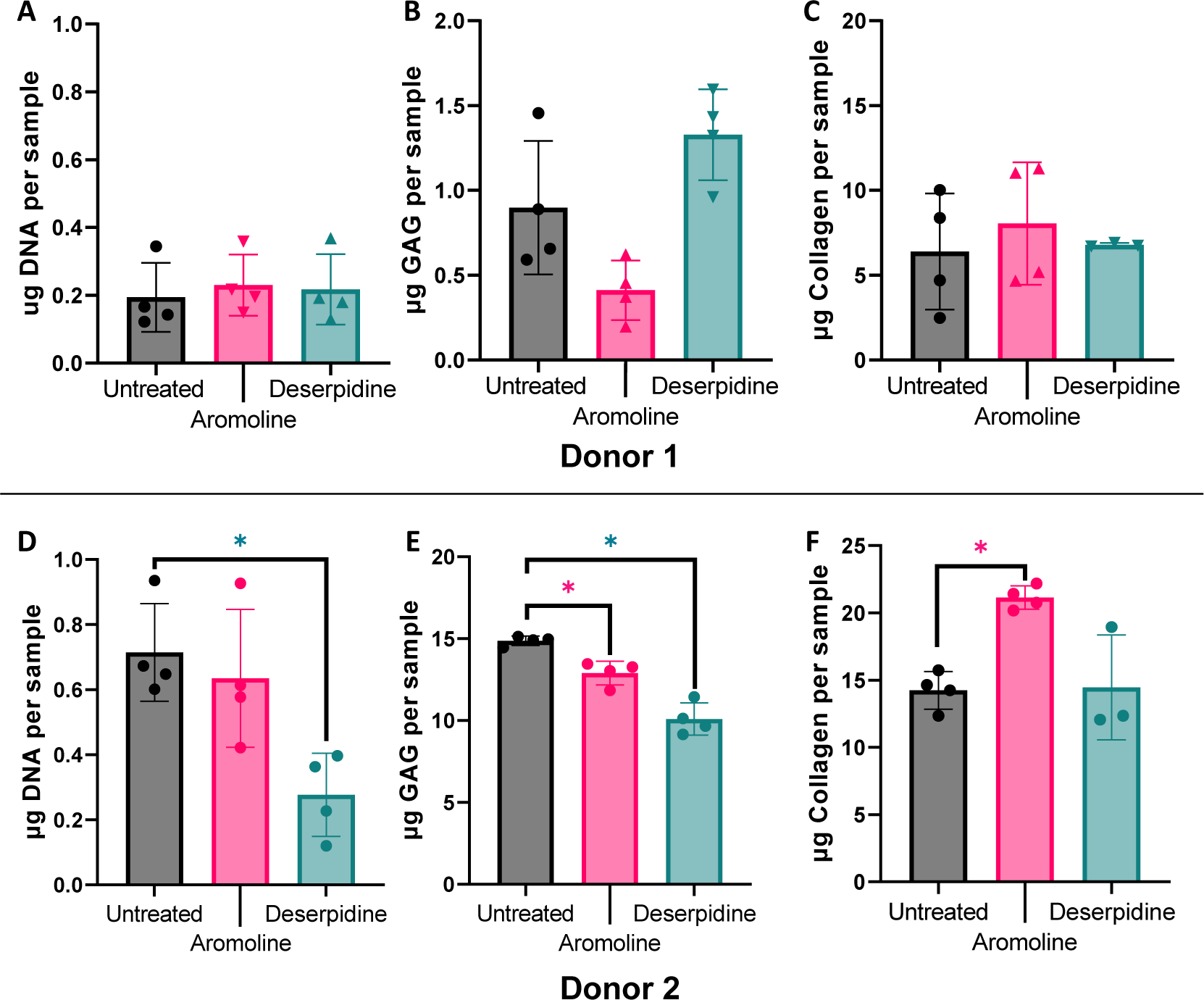
Biochemical assays on HuCol2gLuc aggregates from two donors at day 22. Total DNA content for donor 1 (μg per sample) is shown in (**A**) and for donor 2 in **(D)** Glycosaminoglycan content was quantified in (**B**) for donor 1 and (**E**) for donor 2. Total of micrograms of collagen content for donor 1 and donor 2 are shown in (**C**) and (**F**) respectively. N = 4. Individual replicates or mean of replicates are shown with error bars indicating standard deviation and * indicating p <0.01 vs. untreated control, * aromoline; * deserpidine.

### Aromoline has a dose dependent effect on HuCol2gLuc aggregate type II collagen expression

To characterize the response of HuCol2gLuc cells to treatment with aromoline, cells were cultured in 3D aggregates in chondrogenic media treated with different concentrations of aromoline (0-10 μM). Dose response curves were generated from luminescence data at day 3 (**Fig. 4A**), day 10 (**Fig. 4B**) and for cumulative luminescence (**Fig. 4C**). As seen in Figure 4, there was a dose dependent increase in type II collagen driven-luminescence with a calculated 50% effective concentration (EC50) of 2.46 μM, 2.58 μM, and 3.07 μM for day 3, 10 and cumulative luminescence respectively. To determine effects of aromoline on viability and cell proliferation, resazurin assay and quantification of DNA content were carried out at day 22. **Fig. 4D** shows that there is a dose dependent decrease in resazurin fluorescence at day 22 suggestive of decreased metabolic activity due to aromoline treatment. DNA quantification shows a similar trend with a significant decrease in aggregates treated with 10 μM aromoline (**Fig. 4E**). This suggests that decreased metabolic activity is due to cytotoxicity at 10 µM aromoline. Immunohistochemistry of sections collected from aggregates at day 22 shown in **Fig. 4F-4I** confirm expression of type II collagen in HuCol2gLuc derived aggregates with no notable differences between aggregates treated with these doses of aromoline.

**Fig. 4.**
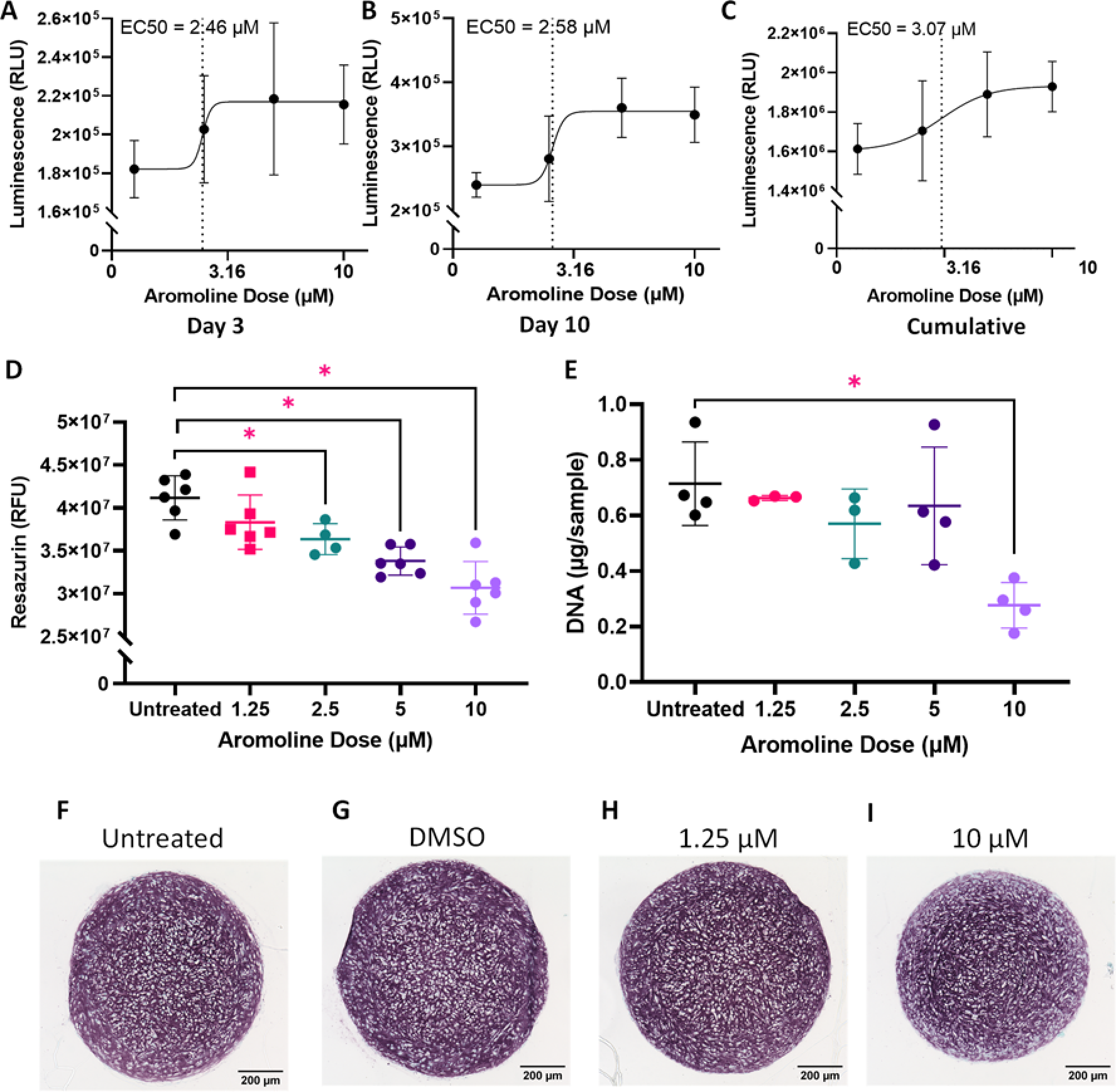
Response of HuCol2gLuc aggregates to aromoline is dose dependent. HuCol2gLuc aggregates were treated with aromoline (0-10 μM). Dose response curves were generated from luminescence data at day 3 (**A**), day 10 (**B**) and cumulative luminescence over 22 days (**C**). Metabolic activity (**D**) and DNA content (**E**) are shown at day 22. N= 6. Individual replicates or mean of replicates are shown with error bars indicating standard deviation and * indicating p <0.05 vs. untreated control. Type II collagen staining of day 22 aggregates treated with indicated doses of aromoline (**F-I**). Scale Bars = 200 µm.

### Characterization and target identification of aromoline

We attempted to identify potential targets for aromoline, by using its chemical structure **(Fig. 5A)** for *in silico* analysis via SwissADME and SwissTargetPrediction ^24–26^. Using the Swiss target prediction tool, we determined that about 40% of the predicted targets are family A G-protein coupled receptors (**Fig. 5B**). By homology, the specific target with the highest probability was the Dopamine 2 receptor (*DRD2*). Interestingly, several of the other compounds from the screen also showed dopamine receptors as a target (Supplemental Data 1a). Our previous work showed expression of only the dopamine 4 receptor (*DRD4*), a DRD2-like receptor, in human chondrocytes ^27^. Another study looking at mesenchymal stromal cell chondrogenesis, via RNA-Seq, also showed DRD4 as the only dopamine receptor expressed during chondrogenesis (**Fig. 5C**; ^28^). Interestingly, expression of DRD4 increases during chondrogenesis with a peak at around week 2 (**Fig. 5C**), similar to the type II collagen pattern observed for aggregates treated with aromoline (**Fig. 2C**). Furthermore, immunohistochemistry confirmed the expression of DRD4 in tissue engineered human cartilage (**Fig. 5D**), with a different more punctate pattern of staining as compared to DRD4 staining in the mouse brain where dopamine receptors are widely expressed.

**Fig. 5.**
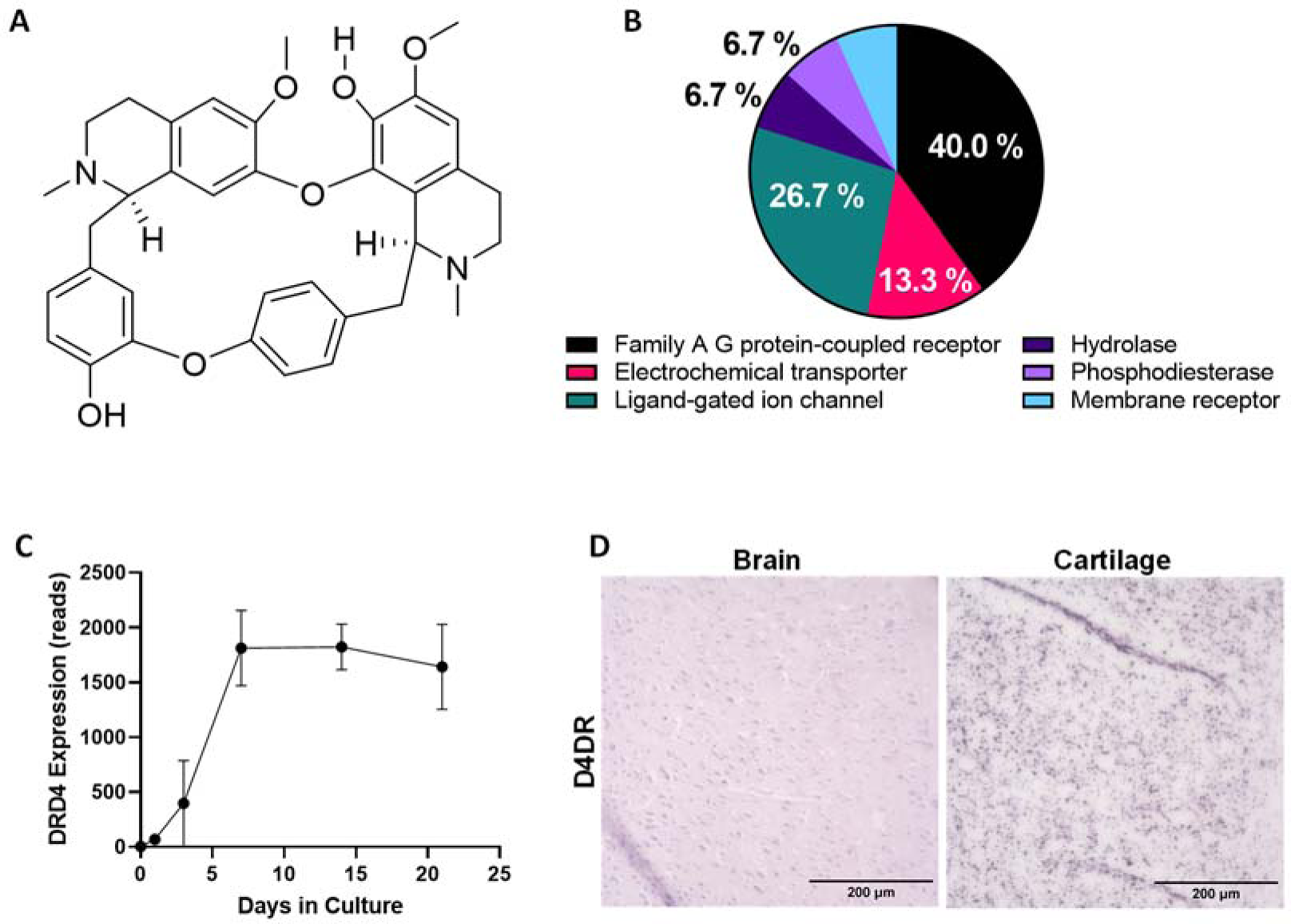
Structure and predicted targets for aromoline. Chemical structure of aromoline (**A**). SwissTargetPrediction predicted targets for aromoline presented by target class (**B**). DRD4 temporal expression in human mesenchymal stromal cells (MSCs) during 21 days of chondrogenesis ((28) **C**, n = 3). DRD4 staining (purple) in mouse brain and in tissue engineered human cartilage (**D**) Scale Bars = 200 µm.

### DRD4 selective antagonist and agonists have a dose dependent effect on HuCol2gLuc aggregate type II collagen expression

To determine whether DRD4 signaling affected type II collagen expression in human chondrocytes, HuCol2gLuc cells from donor 2 were cultured in 3D aggregates in the presence of a selective DRD4 agonist (ABT 724) or antagonist (PNU 96415E) at various concentrations (0-25 μM) and luminescence assessed over 22 days. As seen in **Fig. 6A**, ABT 724 treatment resulted in an increase in luminescence at day 8 at 10 µM and 25 µM concentrations with a sustained signal until day 22. ABT 724 had no effect at 0.1 µM or 1 µM. Cumulative luminescence (**Fig. 6B**) corroborates the results with concentrations of 10 µM and 25 µM resulting in a significant increase in luminescence as compared to the untreated controls. Unexpectedly, treatment with antagonist, PNU 96415E (**Fig. 6C** and **6D**) also had a positive effect on type II collagen-driven luciferase expression with the higher concentrations, 12.5 μM and 25 μM, resulting in increased luminescence compared to the untreated controls. The pattern of expression over 22 days, seen in **Fig. 6C**, differed from treatment with ABT 724 (**Fig. 6B**), with PNU 96415E treatment resulting in peak expression by day 10 followed by a decrease at day 15 and sustained expression until day 22. Cumulative luminescence (**Fig. 6D**) corroborates the results with concentrations of 12.5 μM and 25 μM resulting in a significant increase in luminescence as compared to the untreated controls. Immunohistochemistry confirmed the expression of type II collagen in aggregates treated with ABT 724 and PNU 96415E. ABT 724 had more intense staining for type II collagen as compared to the untreated control, while PNU 96415E had diminished type II collagen staining as compared to the control. DRD4 immunohistochemistry similarly shows a substantial increase in staining for aggregates treated with ABT 724 as compared to the control (**Fig. 6E**). Whereas PNU 96415E had similar staining as compared to the untreated control (**Fig. 6E**).

**Fig. 6.**
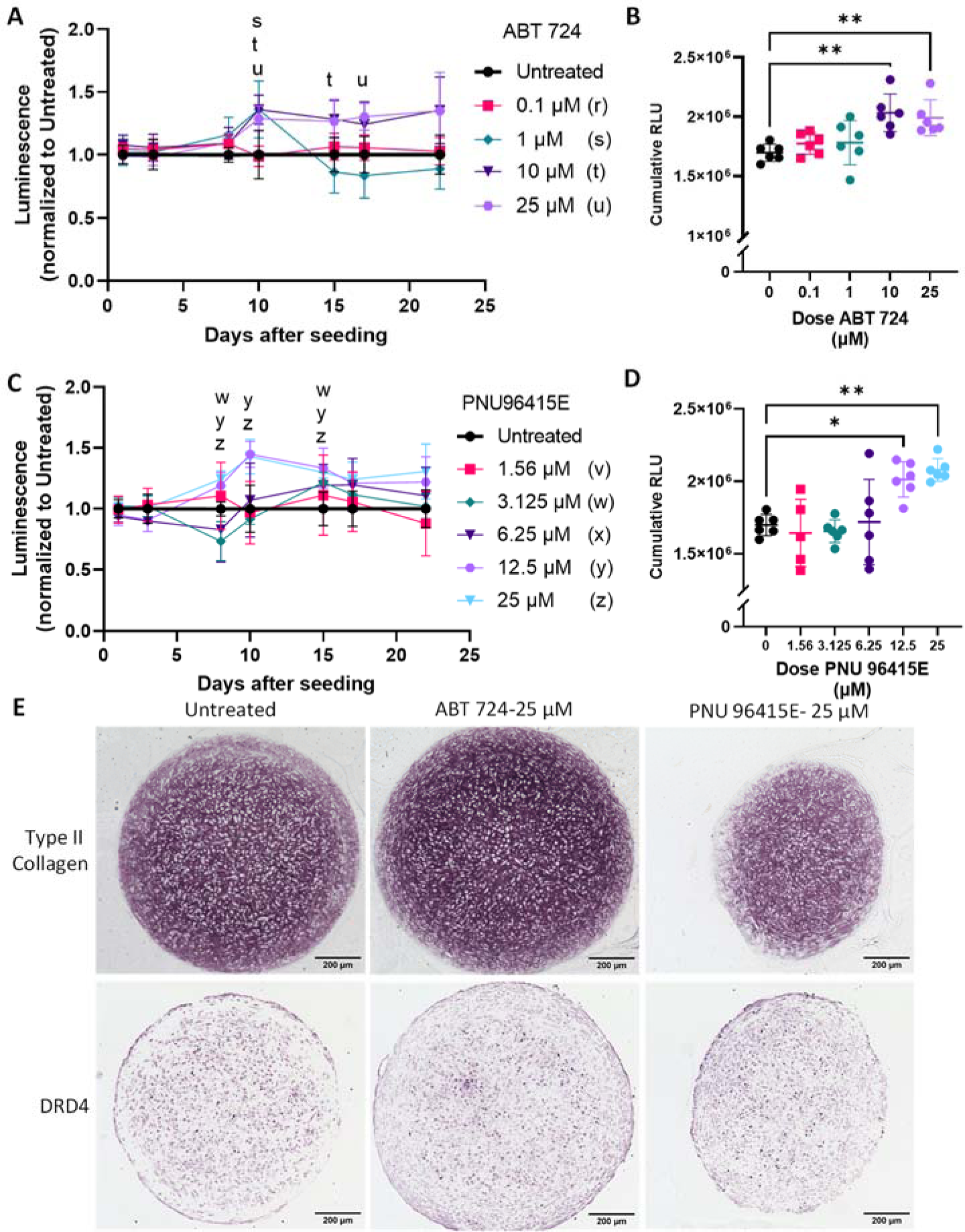
DRD4 selective agonist and antagonist treatment increased type II collagen expression in HuCol2gLuc aggregates. Luminescence signal over 22 days after treatment with DRD4 agonist (ABT 724 (**A**)) and DRD4 antagonist (PNU 96415E (**B**)). Statistically significant differences vs. untreated control are indicated by the respective letters for the dose (s, t, u, w, y, z). Results also shown as cumulative luminescence for treatment with ABT 724 (**C**), and PNU 96415E (**D**). N = 6. Individual replicates or mean of replicates are shown with error bars indicating standard deviation and * indicating p <0.05, and ** indicating p <0.01 vs. untreated control. Immunohistological staining for DRD4 and type II collagen in HuCol2gLuc aggregates after ABT 724 and PNU 96415E treatment at indicated doses **(E)** Scale Bars = 200µm.

To assess whether aromoline had an effect on DRD4 expression in human chondrocytes, monolayer culture of human chondrocytes from donor 1 and donor 2 were treated with drug aromoline (5μM) for 24 h followed by RNA extraction and analysis via qPCR. As seen in **Fig. 7A and 7B**, treatment with aromoline resulted in significant upregulation of DRD4 in both donor 1 and donor 2. Donor 1 had overall higher DRD4 expression as compared to the untreated control, approximately 5-fold more, while donor 2 had only about 1.8-fold higher expression. Histology of donor 2 HuCol2gLuc aggregates treated with aromoline (10 μM) at day 22 shown in **Fig. 7C** confirm increased expression of DRD4 expression compared to untreated controls.

**Fig. 7.**
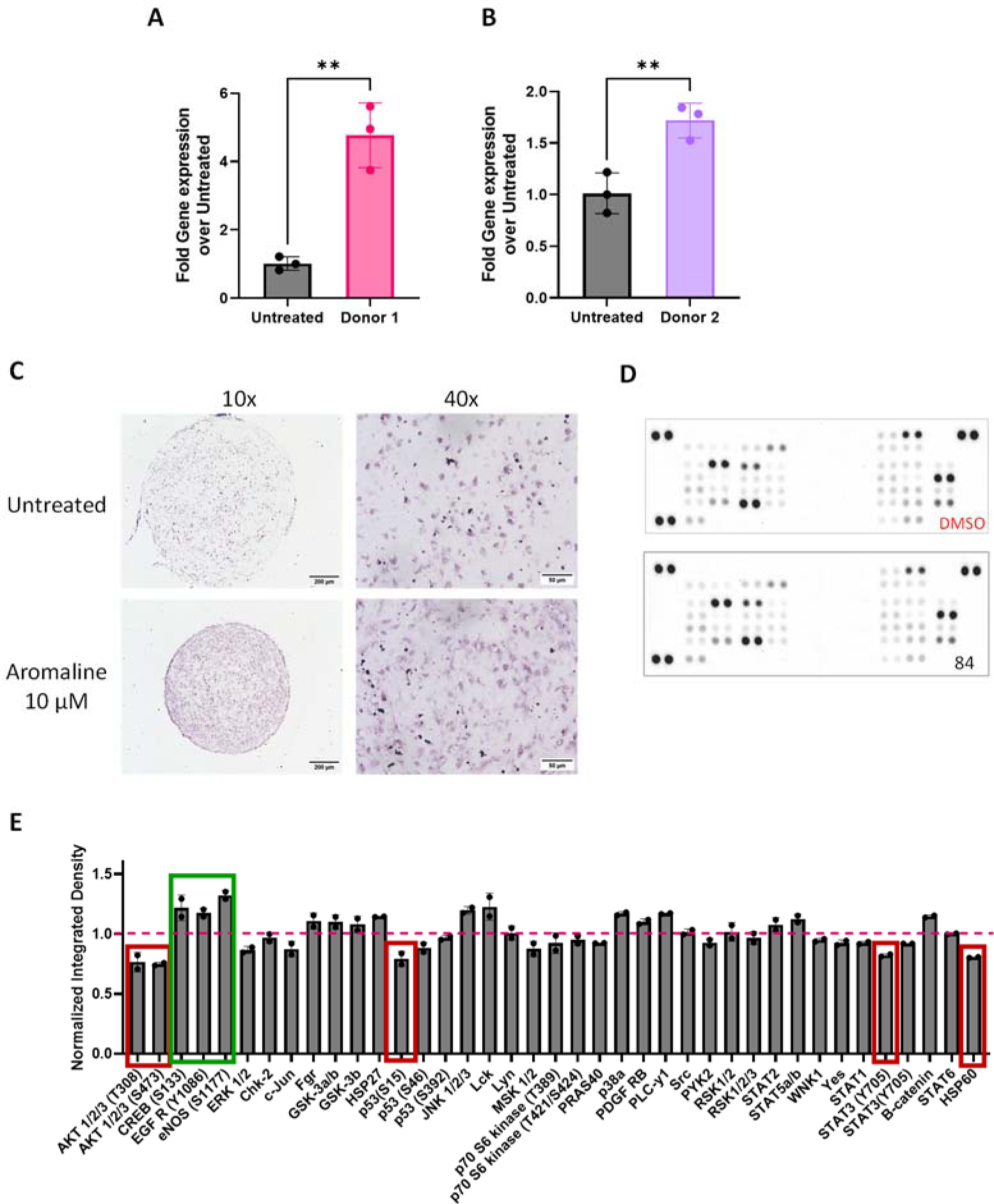
Aromoline increases DRD4 expression in primary human chondrocytes. Gene expression of *DRD4* in monolayer cultures of primary human chondrocytes via RT-qPCR after treatment with aromoline for 24 hours: donor 1 (**A**) and donor 2 (**B**). Results shown as fold change vs. untreated control and normalized to reference gene *HPRT*. N = 3. Error bars indicate standard deviation and ** indicate p <0.01. Immunohistological staining (purple) for DRD4 in HuCol2gLuc aggregates after treatment with aromoline (**C**). Scale Bars = 200 µm. Dot blot of kinase phosphorylation of primary human chondrocytes after treatment with aromoline or DMSO as a vehicle control (**D**). Data from the dot blots is also shown as normalized integrated density (**E**).

To determine what signaling pathways could be involved in aromoline mediated increase of type II collagen, donor 2 primary chondrocytes were treated with aromoline in monolayer culture for 24 h, lysate collected, and a phospho-kinase array was used to look at levels of phosphorylation for 17 different kinases (**Fig. 7D**). Data normalized to reference spots and to DMSO controls is shown in **Fig. 7E**. There is a substantial decrease in AKT 1/2/3 phosphorylation at both T308 and S473 phosphorylation sites (**Fig.7E**). A less considerable decrease in p53 phosphorylation at the S15 site, in STAT3 phosphorylation at Y705, and in HSP 60 was also observed (**Fig. 7E**). Also observed was a modest increase in phosphorylation of CREB, EGFR, and eNOS (**Fig. 7E**).

## DISCUSSION AND CONCLUSIONS

Using our high throughput model of COL2A1-*Gaussia* luciferase primary human chondrocyte reporter (HuCol2gLuc) cells in 3D aggregates under physioxia, we have successfully screened an NCI Natural product library. Our system has several advantages over other proposed models. It makes use of human primary cells over non-human cells or immortalized cell lines. While primary chondrocytes are generally limited in availability and could be a barrier for high throughput screening, our use of porcine derived synoviocyte matrix for expansion preserves chondrogenic capacity for more cell divisions and allowed for greater expansion of primary chondrocytes ^29, 30^. Our system tests compounds in 3D aggregates while other chondrogenic models for drug discovery typically rely on 2D culture ^31–33^. Two-dimensional culture for many cell types, including chondrocytes, fail to mimic cell to cell and cell to matrix interactions, as well as paracrine signaling events that are responsible for physiological tissue structure ^34, 35^. Furthermore, 2D culture cannot correctly model events, such as ECM sequestering of soluble compounds, which can have a large effect on drug - cell interactions ^35–37^. Overall 3D models are more likely to replicate the complexities and responses of native tissue. The primary advantage of our reporter model is the use of a secreted reporter which not only allows us to quantitate the signal as opposed to other read out methods such as staining and microscopy imaging, but also allows for temporal non-destructive assessment of the phenotype during chondrogenesis ^10^.

From the natural product screen, we identified several candidate compounds that increased type II collagen over controls at one or more timepoints (Supplemental Data 1b). Of these, aromoline and deserpidine had the most pronounced effect. In a follow up experiment aromoline presented as the better candidate while deserpidine had inconclusive results. Aromoline (NSC93674) is a bisbenzylisoquinoline alkaloid generally derived from members of the *Berberis* and *Stephania* genus ^38, 39^. As their name suggests, and as seen in **Fig. 5A**, these alkaloids consist of two benzylisoquinoline parts linked through either diphenyl ether, benzyl phenyl ether, or biphenyl bonds ^39, 40^. Previous screening of alkaloids from *Berberis* have shown that aromoline is a potent inhibitor of butyrylcholinesterase with low blood brain barrier permeability ^41^. However, neither our RNA-Seq data nor that of Huynh et al. showed expression of butyrylcholinesterase in chondrocytes or their progenitors ^27, 28^. Another study showed weak anti-microbial activity against *Plasmodium falciparum* with low cytotoxicity in the KB cell line ^42^. Although there is little published research on aromoline, various bisbenzylisoquinoline alkaloids are being studied for their anti-proliferative, anti-inflammatory and anti-microbial properties ^43–48^. This is the first study to our knowledge that explores the effects of aromoline on chondrocytes and chondrogenesis.

With so little known about the signaling pathways activated by aromoline, *in silico* analysis provided an unexpected initial candidate target: the dopamine 2 receptor. Dopamine receptors are G-coupled receptors widely expressed through the central nervous system ^49, 50^. Expression in the periphery has been observed in immune cells such as neutrophils, basophils, B cells and K cells, as well as the heart, kidneys, adrenal glands, blood vessels, and gastrointestinal tract ^51–59^. Dopamine receptors are classified under two groups, the D1-like family that is coupled to G_s_ and activates adenylyl cyclase and the D2-like family that is coupled to G_iα_ and inhibits adenyl cyclase ^60–62^. Based on the *in silico* target analysis, we used a novel approach combining the *in silico* information with transcriptional data. We used our previous RNA-seq data and that of Huynh et al. (**Fig. 5C**), combined with the fact that dopamine receptors from the same family share significant homology, we hypothesized that Dopamine 4 Receptor (DRD4), one of the members of the D2-like receptors was a potential target for aromoline ^27, 28, 50^. This is the first study to our knowledge that has highlighted the expression of DRD4, by RNA-Seq, qPCR and by immunohistochemistry, in primary human chondrocytes and cartilage. Furthermore, we confirmed upregulation in expression of DRD4 and type II collagen after treatment with aromoline (**Fig 7A-7D**).

Interestingly, treatment of HuCol2gLuc aggregates with either a selective agonist or antagonist for DRD4 resulted in increased type II collagen promoter-driven luminescence. However, the antagonist displayed a detrimental effect on pellet size, glycosaminoglycan content (**Supplemental Fig. 1**), and a decreased intensity in type II collagen histology staining (**Fig. 6E**). This suggests that the luminescence results for antagonist treatment are potentially false positives, a common and accepted aspect of screens, and highlight the importance of confirmation via alternative methods.

The best characterized pathway for extracellular matrix deposition by chondrocytes or their progenitors is that of transforming growth factor-beta superfamily and downstream effectors ^63–65^. In the non-canonical pathway, PI-3 kinase is stimulated, phosphorylating Akt which goes on to phosphorylate mTOR, resulting in metabolic changes including protein expression. Aromoline treatment in chondrocytes results in both upregulation of DRD4 expression and decrease in phosphorylation of Akt (**Fig. 7**), a distinct mode of action from both the canonical and non-canonical TGFβ pathways. A dopamine receptor G_iα_ independent pathway, the β-arrestin 2 pathway, could account for the observed effects of aromoline. The β-arrestin 2 pathway is activated as a mechanism for dopamine receptor internalization and desensitization ^66^. Formation of this complex results in inactivation of Akt by protein phosphatase 2 and activation of glycogen synthase kinase-3 (GSK3) signaling ^67^. GSK3 has been linked to maintenance of the chondrocyte phenotype. Inhibition of GSK3 results in cartilage destruction and progression of chondrocytes to terminal differentiation ^68–71^. Previous studies on dopamine receptors, as well as other GPCRs, have shown that ligand activation of β-arrestin 2 is distinct from its G protein dependent activity ^72–76^. Alternatively, the decrease in STAT3 phosphorylation and increase in CREB phosphorylation could be mechanisms for further research ^77^. These observations provide clues as to the mechanism of action for aromoline but fail to completely elucidate the pathways involved. As the production and deposition of extracellular matrix is both temporally and mechanistically distant from receptor activity, this is perhaps unsurprising. Further studies to unravel the signaling pathways involved in aromoline mediated type II collagen expression as well as DRD4 involvement in chondrogenesis are needed. Other areas of study include the identification of the endogenous ligand for DRD4 in chondrocytes and its role in chondrogenesis and osteoarthritis.

The effects of aromoline between donors points to the possibility of donor dependent effects. While a similar trend, increasing the expression of type II collagen prior to the plateau phase around day 10, was evident with all donors, there was a significant difference in the extent of their response. This could be an interesting avenue for further, patient specific, tailored care. What is encouraging is the expression of DRD4 in cartilage in three separate studies: this research, the previous RNA-Seq data from our group ^27^ and in the work by Huynh et al. ^28^. There are multiple steps that need to be completed before translation to the clinic, confirmation in more donor chondrocytes, with non-modified chondrocytes and in an animal model(s).

This study not only provides novel insights into the complex process of chondrogenesis and extracellular matrix deposition but identifies a potential new target receptor and drug candidate for the anabolic treatment or prevention of osteoarthritis.

## EXPERIMENTAL PROCEDURES

To identify novel, chondrogenesis promoting drugs, human primary chondrocytes were genetically modified to express a secreted luciferase reporter under the control of the type II collagen promoter. Those cells were then used in a 3D culture system under physiological oxygen tension to screen a natural product library for promotion of the articular cartilage phenotypic marker, type II collagen. Identified compounds were further screened using in silico reference data to identify potential drug targets. The dopamine receptor D4 was identified and investigated as a potential target with significant effects on type II collagen expression.

### Human primary chondrocyte isolation

Articular cartilage was isolated from visually intact areas of discarded surgical tissue as previously described ^13^. Tissue identified as donor 1 and donor 2 was collected from two donors during total joint re-placement surgery with informed consent under IRB approved protocols (Baylor College of Medicine, H-36683, H-36374). Tissue identified as donor 3 was acquired from a male neonate (4-days old at time of death with alobar holoprosencephaly) through the International Institute for the Advancement of Medicine (IIAM). Tissue (< 3h post-operation or <24h post mortem on ice in saline) was briefly stored in defined chondrogenic media (∼4h, room temperature) before cartilage was dissected under sterile conditions. Isolated cartilage was diced into <1mm^3^ pieces before digestion, first in hyaluronidase for 30 min (660 Units/ml Sigma, H3506; in DMEM/F12, 40ml), then by collagenase type II for ∼16 hours at 37°C (583 Units/ml Worthington Biochemical Corp.; in DMEM/F12 with 10% FBS, 40ml). The digest was filtered through a 70 µm cell strainer, washed once with DMEM/F12, and resuspended in growth media (DMEM/F12 supplemented with 10% FBS (mesenchymal stromal cell selected ^78^), 1% pen/strep). Cells were subsequently infected as described below or cryopreserved (95% FBS, 5% DMSO).

### Lentiviral construct

Lentivirus was generated as previously described ^10, 13^. Briefly, custom COL2A1-*Gaussia* Luciferase plasmid (HPRM22364-LvPG02, GeneCopoeia, Inc.), envelope (pMD2.G) and packaging (psPAX2) plasmids were amplified in *Escherichia coli* (GCI-L3, GeneCopoeia) and purified via silica column (Qiagen Maxiprep) before co-transfection into HEK293Ta (GeneCopoeia) cells via calcium chloride precipitation. Newly packaged lentiviral particles were collected in culture medium after 48h and concentrated via ultracentrifugation (10,000 RCF, 4°C, overnight). Titers for COL2A1-Gluc lentivirus were estimated via real-time PCR and aliquots stored at -80°C.

### Lentivirus infection of primary Human Chondrocytes

Isolated primary human chondrocytes, from each donor, were seeded at 6,100 cells/cm^2^ in growth media (DMEM and allowed to adhere overnight (∼20% confluency). Cells were infected with lentivirus (COL2A1-GLuc; MOI 25 in growth media) in the presence of 4 µg/ml polybrene (Opti-mem, Gibco) for 15 minutes at 4°C followed by overnight incubation at 37°C. Lentiviral medium was replaced with growth medium, and cells expanded to ∼70-90% confluency. Cells were trypsinized (trypsin/EDTA 0.25%), then seeded on synoviocyte matrix coated flasks ^29^. Primary chondrocyte infection was done in physioxic (37°C, 5% O_2_, 5% CO_2_) conditions ^13, 79^. Newly generated COL2A1-GLuc cells were cryopreserved at the end of this first passage (95% FBS, 5% DMSO). These cells were used for all subsequent studies.

### Chondrogenic Culture

Human COL2A1-GLuc or uninfected cells were thawed and seeded in growth media on synoviocyte matrix at 6,000 cells/cm^2^ and expanded to 90-100% confluence in physioxia ^79, 80^. Cells were trypsinized (0.25% Trypsin/EDTA; Corning), resuspended in growth media (for monolayer culture) or chondrogenic differentiation media for 3D aggregates (93.24% High-Glucose DMEM (Gibco), 1% dexamethasone 10^-5^M (Sigma), 1% ITS+premix (Becton-Dickinson), 1% Glutamax (Hyclone), 1% 100 mM Sodium Pyruvate (Hyclone), 1% MEM Non-Essential Amino Acids (Hyclone), 0.26% L-Ascorbic Acid Phosphate 50mM (Wako), 0.5% Fungizone (Life Technologies) with TGFβ1 (Peprotech) and seeded as described below.

### Generation and Maintenance of 3D aggregates

To generate 3D aggregates, cells were seeded at 50,000 cells per well (in 96-well cell repellent u-bottom plates, GreinerBio) and then centrifuged at 500 RCF, 5 min. For the initial drug screen, aggregates were cultured in basal chondrogenic media (1ng/ml TGFβ1) alone, with DMSO (vehicle control) or 5 μM of the drug compounds from the NCI library (Natural Products V Set, (>90% by ELSD, major peak has the correct mass ion) NCI). For candidate validation and subsequent studies aggregates were cultured in basal chondrogenic media (10ng/ml TGFβ1) alone, with DMSO (vehicle control) or compounds as indicated in the results. DRD4 selective agonist ABT 724 and selective antagonist PNU 9641E were used at 1-25 µM based on prior literature^81–83^. Aggregates were cultured for three weeks in physioxic conditions, culture medium was sampled and replaced three times a week with fresh medium. An OT-2 (Opentrons) python coded robotic pipette, programmed at an aspiration height of 2mm from the bottom of the wells and aspiration rate of 40 μl/s was utilized for addition of compound, cell feeding, and media sampling for luciferase assay^13^. After three weeks, cell aggregates were either fixed in neutral buffered formalin for histology or medium removed and aggregates frozen (-20°C) for biochemical assays.

### Luciferase Assay

Conditioned culture medium sampled from aggregates in 96-wells (20µL/well) was assessed using a stabilized *Gaussia* Luciferase buffer mix (50 μl/well) for a final concentration of 0.09 M MES, 0.15M Ascorbic Acid, and 4.2µM Coelenterazine in white 96-well plates. Luminescence was measured in a plate reader (25°C, relative light units (RLU), EnVision plate reader). An OT-2 (Opentrons) python coded robotic pipette was utilized for luciferase buffer addition to white plates (GreinerBio).

### Metabolic Assay (Resazurin)

Metabolic activity was assessed on day 22 by adding resazurin (TCI chemicals) to a final concentration of 50 µM to each well and incubating at 37 °C in physioxia for 3 hours. After three hours, media (120 µl) was transferred to a 96-well black plate (Greinerbio) and fluorescence read at an excitation of 535nm and emission at 588nm. An OT-2 (Opentrons) python-coded robotic pipette was used to add resazurin to cell plates as well as to transfer medium to black plates.

### Immunohistochemistry

At the end of three-week culture, aggregates were fixed in 10% Neutral Buffered Formalin overnight, and subsequently embedded in paraffin wax and sectioned (7µm sections). Sections were deparaffinized and hydrated, followed by treatment with pronase (1mg/ml, Sigma P5147, in PBS with 5mM CaCl_2_) for epitope retrieval. Sections were blocked (BSA 3% w/v, Cohn Fraction V, Alfa Aesar) and incubated with mouse primary anti-Collagen Type II (DSHB II-II6B3) or mouse anti-dopamine 4 receptor (MABN125, Millipore) primary for 2 h at room temperature in a humidified chamber. II-II6B3 was deposited to the DSHB by Linsenmayer, T.F. (DSHB Hybridoma Product II-II6B3). Sections were then incubated with a biotinylated secondary (Vector Labs) followed Streptavidin-HRP (BD Biosciences). Sections were stained with a chromogen-based substrate kit (Vector labs, VIP substrate vector kit). All sections were imaged using a Keyence BZ-X810 microscope at 10x magnification.

### Biochemical Assays

Aggregates frozen at day 22, were thawed with PBS, and digested with papain (25 µg/ml, Sigma, P4762, in 2mM cysteine; 50mM sodium phosphate; 2mM EDTA at a pH 6.5, 100 µl) at 65°C overnight ^13, 79, 84^. Plates were sealed with a qPCR adhesive sealing film (USA Scientific), a silicone sheet, and steel plates clamped to the plate to prevent evaporation during digestion. Half of the digest (50 μl) was transferred to another plate and frozen for hydroxyproline assessment. In the remaining digest, papain was inactivated with 0.1M sodium hydroxide (NaOH, 50 µl) followed by NaOH neutralization (100mM Na_2_HPO_4_, 0.1 N HCL, pH 1.82, 50 µl). To quantitate DNA content, samples of the neutralized digest (20 µl) were combined with buffered Hoechst dye (#33258, 667ng/ml, phosphate buffer pH 8, 100 µl) and fluorescence measured at an excitation of 365nm and emission of 460nm. For GAG assessment, 1,9-Dimethyl-methylene blue solution (195 µl) was added to neutralized digests (5 µl) and absorbance measured at 595nm and 525nm ^85^. Readings were corrected by subtracting 595nm reading from 525nm. Micrograms of DNA and GAG were calculated using a Calf thymus DNA standard (Sigma) or Chondroitin Sulfate standard (Seikagaku Corp.), respectively.

Hydroxyproline (HP) was quantified as previously described^86^. The frozen digests (50 μl) were thawed at room temperature and incubated overnight at 105°C with 6M hydrochloric acid (200 μl). Plates were sealed to prevent evaporation. Samples were then dried at 70°C overnight with a hydroxyproline standard (Sigma). Copper sulfate (0.15M, 10µl) and NaOH (2.5M, 10µl) were added to each well and incubated at 50°C for 5 minutes, followed by hydrogen peroxide (6%, 10 µl) for 10 minutes. Sulfuric acid (1.5 M, 40 μl) and Ehrlich’s reagent (20 µl) were added, and samples incubated at 70°C for 15 minutes ^79^. Absorbance was measured at 505nm. Micrograms of hydroxyproline were calculated using the standard. Total collagen was calculated by the following formula (µg of HP X 7.6 = µg Total Collagen).

### Reverse Transcription and Real-Time PCR

Primary human chondrocytes were seeded at 280,000 cells/well (12-well adherent plate, Corning) in growth media and allowed to adhere overnight. Growth media was replaced with chondrogenic media supplemented with 1ng/mL TGFβ1 alone or with aromoline. Cells were treated for 24 h and then lysed and frozen. Lysate was thawed and RNA extraction and purification done following manufacturer’s protocol (Ambion PureLink RNA Mini kit). RNA purity and integrity was assessed by RNA ScreenTape (Agilent Technologies) before use. cDNA was synthesized from 400ng RNA using a Maxima H Minus reverse transcriptase master mix following manufacturer’s protocol. Quantitative real-time PCR for *DRD4* and *HPRT* (endogenous control) gene expression was done (qPCR) using SYBR green master mix (Applied Biosystems) and QuantStudio7 Flex Real-Time PCR system (ThermoFisher Scientific). Cycling parameters: 95°C for 20s then 45 cycles of 95°C 10s, 60°C 20s, 72°C 19s, followed by melt curve analysis. Primers: *HRPT*, forward primer: 5’ ATTGACACTGGCAAAACAATGC 3’, reverse primer: 5’ TCCAACACTTCGTGGGGTCC 3’, reference. *DRD4*, forward primer: 5’ GTGGTGGTCGGGGCCTT 3’, reverse primer: 5’ CGGAGCAGGCAGGACAC 3’. *DRD4* CT values were normalized to *HRPT* expression and *DRD4* relative gene expression vs untreated calculated.

### Phospho-kinase Array

Primary human chondrocytes were seeded at 5 x 10^6^ cells (2 x 10 cm adherent dish, Corning) in growth media and allowed to adhere overnight. Growth media was replaced with chondrogenic media supplemented with 1ng/mL TGFβ1 with DMSO or with aromoline (5μM). Cells were treated for 24 h before processing according to the manufacturer’s protocol (ARY003C, R&D Systems). This phospho-kinase array contains capture antibodies to measure relative levels of phosphorylation of 37 kinases on a nitrocellulose membrane. Signal intensity was quantified using ImageJ using the Microarray Profile plug-in.

### Statistical Analysis

Statistical analysis for all experiments was performed using GraphPad Prism 9 and volcano plot, multiple t-test with 1% false detection rate, one-way or two-way ANOVA with Dunnett’s correction performed. All data passed tests for normality. In all figures * indicates p-value < 0.05, ** indicates p-value <0.01, and *** indicates p-value < 0.001.

## Acknowledgments

We gratefully acknowledge the donors and the International Institute for the Advancement of Medicine (IIAM) for the tissue used in these studies. The plasmids, pMD2.G, and psPAX2, were a gift from Didier Trono (Addgene plasmid # 12259; http://n2t.net/addgene:12259; RRID:Addgene_12259) and (Addgene plasmid # 12260; http://n2t.net/addgene:12260; RRID:Addgene_12260) respectively. The type II collagen antibody, II-II6B3 was deposited to the DSHB by Linsenmayer, T.F. (DSHB Hybridoma Product II-II6B3).

## Funding

Rolanette and Berdon Lawrence Bone Disease Program of Texas (TJK)

University of Central Florida (TJK)

University of Central Florida College of Medicine (TJK).

## Author contributions

Conceptualization: MAC, TJK. Methodology: MAC, TJK. Software: MAC, RM, TJK. Investigation: MAC, SG, MK, KM, RM, TJK. Visualization: MAC, SG, TJK. Funding acquisition: TJK. Project administration: TJK. Supervision: MAC, TJK. Writing – original draft: MAC. Writing – review & editing: MAC, TJK

## Abbreviations Used

ANOVA: Analysis of Variance
CREB: CAMP Responsive Element Binding Protein
COL2A1-Gluc: type II collagen promoter driven *Gaussia* luciferase
DMEM: Dulbecco’s Modified Eagle’s Medium
DRD4: Dopamine Receptor D4
eNOS: endothelial Nitric Oxide Synthase
FBS: Fetal Bovine Serum
GAG: Glycosaminoglycan
HP: Hydroxyproline
HPRT: Hypoxanthine Phosphoribosyltransferase
HRP: Horseradish Peroxidase
HSP 60: Heat Shock Protein 60
MEM: Minimal Essential Media
OA: Osteoarthritis
PTOA: Post-traumatic Osteoarthritis
qPCR: quantitative PCR
STAT3: Signal Transducer And Activator Of Transcription 3
SYBR: N’,N’-dimethyl-N-[4-[(E)-(3-methyl-1,3-benzothiazol-2-ylidene)methyl]-1-phenylquinolin-1-ium-2-yl]-N-propylpropane-1,3-diamine
TGFβ1: Transforming Growth Factor Beta 1

