## supplemental data 1b for "Disease Modifying Osteoarthritis Drug Discovery Using A Temporal Phenotypic Reporter In 3D Aggregates of Primary Human Chondrocytes"

| Sum of probability | Column Labels | | | | | | | |
|---|---|---|---|---|---|---|---|---|
| Row Labels | 84 | 186 | 204 | 391 | 413 | 416 | 418 | Grand Total |
| 11-beta-hydroxysteroid dehydrogenase 1 | | | | | 0.100578902 | | | 0.100578902 |
| 3-phosphoinositide dependent protein kinase-1 | 0.074564954 | | | | | | | 0.074564954 |
| Acetylcholine receptor; alpha1/beta1/delta/gamma | | | | 0.111501865 | | 0.111501865 | 0.097874534 | 0.320878264 |
| Acetylcholinesterase | 0.074564954 | | | | 1 | | | 1.074564954 |
| Acidic mammalian chitinase | | | | | 0.100578902 | | | 0.100578902 |
| Adenosine A1 receptor | | | 0.082221517 | | 0.239574696 | | | 0.321796213 |
| Adenosine A2a receptor | | | 0.082221517 | | 0.19883376 | | | 0.281055277 |
| Adenosine A2b receptor | | | | | 1 | | | 1 |
| Adenosine A3 receptor | | | 0.082221517 | | 0.149732594 | | | 0.231954111 |
| Adenosine kinase | | 0.06423878 | | | | | | 0.06423878 |
| Adrenergic receptor alpha-2 | | | 0.082221517 | 0.158886034 | | 0.182601417 | 0.213125923 | 0.636834891 |
| Adrenergic receptor beta | 0.074564954 | 0.06423878 | 0.082221517 | 0.111501865 | | 0.111501865 | 0.097874534 | 0.541903515 |
| ALK tyrosine kinase receptor | 0.074564954 | 0.06423878 | 0.082221517 | | 0.100578902 | | | 0.321604153 |
| Alpha-1a adrenergic receptor | 0.074564954 | 0.06423878 | 0.082221517 | | | | | 0.221025251 |
| Alpha-1a adrenergic receptor (by homology) | | | | 0.206265233 | | 0.206265233 | 0.295453305 | 0.707983771 |
| Alpha-1b adrenergic receptor | 0.074564954 | 0.06423878 | 0.082221517 | 0.12730257 | | 0.119403562 | 0.122581769 | 0.590313151 |
| Alpha-1d adrenergic receptor | 0.074564954 | 0.06423878 | 0.082221517 | 0.12730257 | | 0.135202128 | 0.155528102 | 0.63905805 |
| Alpha-2a adrenergic receptor | | | 0.082221517 | 0.158886034 | | 0.182601417 | 0.213125923 | 0.636834891 |
| Alpha-2b adrenergic receptor | | | 0.082221517 | 0.158886034 | | 0.182601417 | 0.213125923 | 0.636834891 |
| Amine oxidase, copper containing | | | | 0 | | 0 | 0.097874534 | 0.097874534 |
| Anti-estrogen binding site (AEBS) | | | | 0.111501865 | | 0.111501865 | 0.097874534 | 0.320878264 |
| Apoptosis regulator Bcl-2 | | 0.06423878 | | | | | | 0.06423878 |
| Apoptosis regulator Bcl-X | | 0.06423878 | | | | | | 0.06423878 |
| Arachidonate 12-lipoxygenase | | | | 0.111501865 | | 0.111501865 | 0.097874534 | 0.320878264 |
| Arachidonate 15-lipoxygenase | | | | 0.111501865 | | 0.111501865 | 0.097874534 | 0.320878264 |
| ATP-sensitive inward rectifier potassium channel 1 | | | 0.082221517 | | | | | 0.082221517 |
| Baculoviral IAP repeat-containing protein 2 | 0.074564954 | 0.06423878 | 0.082221517 | | | | 0.097874534 | 0.318899785 |
| Baculoviral IAP repeat-containing protein 3 | 0.074564954 | 0.06423878 | | | | | | 0.138803734 |
| Beta secretase 2 | | 0.06423878 | | | | | | 0.06423878 |
| Beta-1 adrenergic receptor | 0.074564954 | 0.06423878 | 0.082221517 | 0.143102156 | | 0.135202128 | 0.130791955 | 0.63012149 |
| Beta-3 adrenergic receptor | 0.074564954 | 0.06423878 | 0.082221517 | | | | 0.097874534 | 0.318899785 |
| Beta-secretase 1 | | 0.06423878 | | | | | | 0.06423878 |
| Bradykinin B1 receptor | | 0.06423878 | 0.082221517 | | | | | 0.146460297 |
| Brain adenylate cyclase 1 | | 0.06423878 | 0.082221517 | | | | | 0.146460297 |
| Bromodomain testis-specific protein | | | | | 0.149732594 | | | 0.149732594 |
| Bromodomain-containing protein 2 | | | | | 0.215238178 | | | 0.215238178 |
| Bromodomain-containing protein 3 | | | | | 0.215238178 | | | 0.215238178 |
| Bromodomain-containing protein 4 | | | | | 0.223449266 | | | 0.223449266 |
| Butyrylcholinesterase | 0.53011145 | | 0.082221517 | | | | | 0.612332967 |
| Calpain 1 | | | | | 0.100578902 | | | 0.100578902 |
| CaM kinase II | | | | 0.111501865 | | 0 | | 0.111501865 |
| Cannabinoid receptor 1 | | | | | 0.100578902 | | | 0.100578902 |
| Carbonic anhydrase II | | | | | 0.100578902 | | | 0.100578902 |
| Carbonic anhydrase III | | | | | | 0 | | 0 |
| Carbonic anhydrase IV | | | | | | 0 | | 0 |
| Carbonic anhydrase VA | | | | | | 0 | | 0 |
| Carbonic anhydrase VB | | | | | | 0 | | 0 |
| Carbonic anhydrase VI | | | | | | 0 | | 0 |
| Carbonic anhydrase XII | | | | 0 | | 0 | | 0 |
| Carbonic anhydrase XIII | | | | | | 0 | | 0 |
| Casein kinase I delta | | | | 0.111501865 | | 0 | | 0.111501865 |
| Casein kinase I epsilon | | | | 0.111501865 | | 0 | | 0.111501865 |
| Caspase-1 | | | 0.082221517 | | | | | 0.082221517 |
| Cathepsin (B and K) | | | 0.082221517 | | 0.100578902 | | | 0.182800419 |
| Cathepsin (V and K) | | | | | 0.100578902 | | | 0.100578902 |
| Cathepsin D | | 0.06423878 | | | | | | 0.06423878 |
| Cathepsin E | | 0.06423878 | 0.082221517 | | | | | 0.146460297 |
| Cathepsin K | | | | | 0.100578902 | | | 0.100578902 |
| Cathepsin L | 0.074564954 | | | | 0.100578902 | | | 0.175143856 |
| Cathepsin S | | | | | 0.100578902 | | | 0.100578902 |
| C-C chemokine receptor type 3 | | 0.06423878 | 0.082221517 | | | | | 0.146460297 |
| C-C chemokine receptor type 4 | 0.074564954 | | | 0 | | 0.111501865 | | 0.186066819 |
| C-C chemokine receptor type 5 | | 0.06423878 | 0.082221517 | | | | | 0.146460297 |
| C-C motif chemokine 2 | | | | 0.111501865 | | 0.111501865 | 0.097874534 | 0.320878264 |
| CDK8/Cyclin C | | | | | 0.100578902 | | | 0.100578902 |
| CDK9/cyclin T1 | 0.074564954 | | | | | | | 0.074564954 |
| Cell division protein kinase 8 | | | | | 0.100578902 | | | 0.100578902 |
| Ceramide glucosyltransferase | | | 0.082221517 | | | | | 0.082221517 |
| Cholecystokinin B receptor | | 0.06423878 | | | | | | 0.06423878 |
| c-Jun N-terminal kinase 1 | | | | 0 | | | 0.097874534 | 0.097874534 |
| Coagulation factor VII/tissue factor | 0.074564954 | | | 0.111501865 | | 0.111501865 | 0.097874534 | 0.395443218 |
| Complement factor D | 0.074564954 | | 0 | | | | | 0.074564954 |
| C-X-C chemokine receptor type 3 | | 0.06423878 | | | | | | 0.06423878 |
| C-X-C chemokine receptor type 4 | 0.074564954 | | | 0 | | | | 0.074564954 |
| C-X-C chemokine receptor type 7 | | | 0.082221517 | | | | | 0.082221517 |
| Cyclin T1 | | 0.06423878 | | | | | | 0.06423878 |
| Cyclin-dependent kinase 1 | 0.074564954 | | | 0.111501865 | | 0 | | 0.186066819 |
| Cyclin-dependent kinase 2 | 0.074564954 | 0.06423878 | | 0.111501865 | | 0.111501865 | | 0.361807464 |
| Cyclin-dependent kinase 2/cyclin A | | 0.06423878 | 0.082221517 | 0.111501865 | | 0.111501865 | 0.097874534 | 0.467338561 |
| Cyclin-dependent kinase 2/cyclin E | | | | 0 | | | | 0 |
| Cyclin-dependent kinase 2/cyclin E1 | 0.074564954 | | | 0 | 0.100578902 | | | 0.175143856 |

| | | | | | | | | |
|---|---|---|---|---|---|---|---|---|
| Cyclin-dependent kinase 4 | | 0.06423878 | | | | | | 0.06423878 |
| Cyclin-dependent kinase 5/CDK5 activator 1 | | | | | 0.100578902 | | | 0.100578902 |
| Cyclin-dependent kinase 9 | | | | | | | 0.097874534 | 0.097874534 |
| Cyclooxygenase-1 | | | | | | | 0.097874534 | 0.097874534 |
| Cytochrome P450 11B1 | | | | | 0.100578902 | | | 0.100578902 |
| Cytochrome P450 11B2 | | | | | 0.100578902 | | | 0.100578902 |
| Cytochrome P450 1A2 | | | | | | | 0.097874534 | 0.097874534 |
| Cytochrome P450 2D6 | | | 0.082221517 | | | 0 | 0.097874534 | 0.180096051 |
| Cytochrome P450 3A4 | | 0.083326526 | | | | | | 0.083326526 |
| Delta opioid receptor | | | 0.082221517 | 0.111501865 | | 0.111501865 | 0.097874534 | 0.403099781 |
| Dipeptidyl peptidase I | | | | | | | 0.097874534 | 0.097874534 |
| Dipeptidyl peptidase IV | 0.074564954 | 0.06423878 | | 0.111501865 | | 0.111501865 | 0.097874534 | 0.459681998 |
| Dipeptidyl peptidase IX | | | | 0.111501865 | | 0.111501865 | 0.097874534 | 0.320878264 |
| DNA topoisomerase II alpha | | | | | 0.100578902 | | | 0.100578902 |
| DNA-dependent protein kinase | | 0.06423878 | 0.082221517 | | | | | 0.146460297 |
| Dopamine D1 receptor | 0.122098887 | | | 0.617022037 | | 0.78136356 | 0.616846761 | 2.137331245 |
| Dopamine D2 receptor | | | 0.082221517 | | 0.100578902 | | | 0.182800419 |
| Dopamine D2 receptor (by homology) | 0.786967746 | | | 0.617022037 | | 0.78136356 | 0.666318839 | 2.851672182 |
| Dopamine D3 receptor | 0.074564954 | 0.06423878 | 0.082221517 | 0.459066154 | 0.100578902 | 0.459066154 | 0.509599763 | 1.749336224 |
| Dopamine D4 receptor | 0.074564954 | | | 0.237888614 | 0.100578902 | 0.206265233 | 0.246150751 | 0.865448454 |
| Dopamine D5 receptor | 0.074564954 | | | 0.19050211 | | 0.19050211 | 0.237885168 | 0.693454342 |
| Dopamine transporter | | | | 0.269486399 | | 0.3326854 | 0.427205275 | 1.029377074 |
| Dopamine transporter (by homology) | 0.844359912 | | | | | | | 0.844359912 |
| Dual-specificity tyrosine-phosphorylation regulated kinase 2 | | | | | | | 0.097874534 | 0.097874534 |
| dUTP pyrophosphatase | | | 0.082221517 | | | | | 0.082221517 |
| Endothelin receptor ET-A | | 0.06423878 | | | | | | 0.06423878 |
| Epidermal growth factor receptor erbB1 | | 0.06423878 | | 0.111501865 | 0.100578902 | 0 | | 0.276319547 |
| Estrogen receptor alpha | | 0.06423878 | | | | | | 0.06423878 |
| Estrogen receptor beta | | 0.06423878 | | | | | | 0.06423878 |
| Fibroblast activation protein alpha | | | | 0.111501865 | | 0.111501865 | 0.097874534 | 0.320878264 |
| Fibroblast growth factor receptor 1 | | 0.06423878 | 0.082221517 | | | | | 0.146460297 |
| Focal adhesion kinase 1 | 0.074564954 | 0.06423878 | | | | | | 0.138803734 |
| Gamma-secretase | | 0.06423878 | | | 0.100578902 | | | 0.164817682 |
| Geranylgeranyl transferase type I | | | | | 0.100578902 | | | 0.100578902 |
| Glucocorticoid receptor | | | | | 0.100578902 | | | 0.100578902 |
| Glutamate NMDA receptor; GRIN1/GRIN2B | | | 0.082221517 | | | | | 0.082221517 |
| Glycine transporter 1 | | | | | 0.100578902 | | | 0.100578902 |
| Glycogen synthase kinase-3 beta | | | 0.082221517 | | | | | 0.082221517 |
| Gonadotropin-releasing hormone receptor | | | 0.082221517 | | | | | 0.082221517 |
| Heat shock protein HSP 90-alpha | | 0.06423878 | | | | | | 0.06423878 |
| Heat shock protein HSP 90-beta | 0.074564954 | 0.06423878 | | | | | | 0.138803734 |
| Hematopoietic prostaglandin D synthase | | | | | | | 0.097874534 | 0.097874534 |
| Hepatocyte growth factor receptor | 0.074564954 | | 0.082221517 | 0 | 0.100578902 | | | 0.257365373 |
| HERG | | 0.06423878 | | 0.214178827 | 0.149732594 | 0.222013735 | 0.180252494 | 0.83041643 |
| Histamine H1 receptor | | 0.06423878 | | | | | | 0.06423878 |
| Histamine H2 receptor | | | | 0.119403562 | | 0.119403562 | 0.213125923 | 0.451933047 |
| Histamine H3 receptor | | 0.06423878 | 0.082221517 | | | | | 0.146460297 |
| Histamine H4 receptor | | | | 0 | 0.100578902 | | 0.097874534 | 0.198453436 |
| Histone deacetylase 1 | 0.074564954 | | | | | | | 0.074564954 |
| Histone-lysine N-methyltransferase, H3 lysine-79 specific | | 0.06423878 | 0.082221517 | | | | | 0.146460297 |
| Histone-lysine N-methyltransferase, H3 lysine-9 specific 3 | 0.074564954 | | | | | | | 0.074564954 |
| HLA-DR antigens-associated invariant chain | | | 0.082221517 | | | | | 0.082221517 |
| Hydroxycarboxylic acid receptor 2 | | | | | 0.100578902 | | | 0.100578902 |
| Indoleamine 2,3-dioxygenase | | | | 0.111501865 | 0.100578902 | 0.111501865 | 0.097874534 | 0.421457166 |
| Inhibitor of apoptosis protein 3 | 0.074564954 | 0.06423878 | 0.082221517 | | | | 0.097874534 | 0.318899785 |
| Inhibitor of NF-kappa-B kinase (IKK) | 0.149129908 | | | | | | | 0.149129908 |
| Inhibitor of nuclear factor kappa B kinase beta subunit | 0.074564954 | 0.06423878 | | 0.111501865 | | 0.111501865 | | 0.361807464 |
| Insulin receptor | | 0.06423878 | 0.082221517 | | | | | 0.146460297 |
| Insulin-like growth factor I receptor | | | | | 0.100578902 | | | 0.100578902 |
| Integrin alpha-IIb/beta-3 | | 0.06423878 | | | | | | 0.06423878 |
| Interleukin-1 receptor-associated kinase 4 | | | 0.082221517 | 0.111501865 | | 0 | | 0.193723382 |
| Isocitrate dehydrogenase [NADP] cytoplasmic | 0.074564954 | | | | | | | 0.074564954 |
| Kappa Opioid receptor | | | | | | | 0.097874534 | 0.097874534 |
| Kinesin-1 heavy chain/ Tyrosine-protein kinase receptor RET | 0.074564954 | 0.06423878 | | | | | | 0.138803734 |
| Legumain | | | | | 0.100578902 | | | 0.100578902 |
| Leucine-rich repeat serine/threonine-protein kinase 2 | | | | 0 | | | 0.097874534 | 0.097874534 |
| Leukotriene A4 hydrolase | | | | 0.111501865 | | 0.111501865 | 0.097874534 | 0.320878264 |
| LIM domain kinase 1 | 0.074564954 | | | | | | | 0.074564954 |
| Lysine-specific demethylase 5B | | | | | 0.100578902 | | | 0.100578902 |
| Macrophage colony stimulating factor receptor | 0.074564954 | | 0.082221517 | | | | | 0.156786471 |
| Macrophage-stimulating protein receptor | | | 0.082221517 | | | | | 0.082221517 |
| Mammalian target of Rapamycin (mTORC1) | | 0.06423878 | 0.082221517 | | | | | 0.146460297 |
| MAP kinase p38 alpha | 0.074564954 | | 0.082221517 | | 0.100578902 | | | 0.257365373 |
| MAP kinase p38 beta | | | 0.082221517 | | | | | 0.082221517 |
| MAP kinase signal-integrating kinase 2 | | 0.06423878 | 0.082221517 | | | | | 0.146460297 |
| MAP kinase-activated protein kinase 2 | 0.074564954 | | | | | | | 0.074564954 |
| Maternal embryonic leucine zipper kinase | 0.074564954 | 0.06423878 | | | | | | 0.138803734 |
| Matrix metalloproteinase 1 | | 0.06423878 | | | 0.100578902 | | | 0.164817682 |
| Matrix metalloproteinase 9 | | | | | 0.100578902 | | | 0.100578902 |
| Melanin-concentrating hormone receptor 1 | | 0.06423878 | | | | | | 0.06423878 |
| Melanocortin receptor 1 | 0.074564954 | 0.06423878 | | | | | | 0.138803734 |
| Melanocortin receptor 3 | 0.074564954 | 0.06423878 | | | | | | 0.138803734 |
| Melanocortin receptor 4 | | 0.06423878 | | | | | | 0.06423878 |

| | | | | | | | | |
|---|---|---|---|---|---|---|---|---|
| Melanocortin receptor 5 | | 0.06423878 | | | | | | 0.06423878 |
| Melatonin receptor 1A | | | | | 0.100578902 | | | 0.100578902 |
| Melatonin receptor 1B | | | | | 0.100578902 | | 0.097874534 | 0.198453436 |
| Methionine aminopeptidase 1 | 0.074564954 | | | | | | | 0.074564954 |
| Mitogen-activated protein kinase kinase kinase 12 | | | 0.082221517 | | | | | 0.082221517 |
| Mitogen-activated protein kinase kinase kinase 7 | | | 0.082221517 | | | | | 0.082221517 |
| Monoamine oxidase A | 0.074564954 | | | 0 | | | 0.097874534 | 0.172439488 |
| Monoamine oxidase B | | | | | 0.100578902 | | | 0.100578902 |
| Mu opioid receptor | 0.074564954 | | 0.658856029 | 0.19050211 | | 0.182601417 | 0.270836923 | 1.377361433 |
| Multidrug and toxin extrusion protein 1 | 0.559324248 | 0.06423878 | 0.082221517 | | | | | 0.705784545 |
| Multidrug and toxin extrusion protein 2 | | 0.06423878 | 0.082221517 | | | | | 0.146460297 |
| Muscarinic acetylcholine receptor M1 | | | | | 0.100578902 | 0 | | 0.100578902 |
| Muscarinic acetylcholine receptor M1 (by homology) | | | | 0.111501865 | | | 0.097874534 | 0.209376399 |
| Muscarinic acetylcholine receptor M2 | | | | | 0.100578902 | 0 | 0.097874534 | 0.198453436 |
| Muscarinic acetylcholine receptor M3 | | 0.06423878 | | | | 0 | 0.097874534 | 0.162113314 |
| Muscarinic acetylcholine receptor M4 | 0.074564954 | | | | | | 0.097874534 | 0.172439488 |
| Myosin light chain kinase, smooth muscle | | | | | | | 0.097874534 | 0.097874534 |
| NADPH oxidase 4 | | | 0.082221517 | | | | | 0.082221517 |
| Nerve growth factor receptor Trk-A | | | | | 0.100578902 | | | 0.100578902 |
| Neurokinin 1 receptor | | | | | 0.100578902 | | | 0.100578902 |
| Neurokinin 2 receptor | | 0.092751834 | | | | | | 0.092751834 |
| Neuronal acetylcholine receptor protein alpha-4 subunit | | | | 0 | | 0 | 0.097874534 | 0.097874534 |
| Neuronal acetylcholine receptor protein alpha-7 subunit | | | | 0.111501865 | | 0.111501865 | | 0.22300373 |
| Neuronal acetylcholine receptor; alpha2/beta4 | 0.273748951 | | | | | | | 0.273748951 |
| Neuronal acetylcholine receptor; alpha3/alpha6/beta2/beta3 | | | | | | | 0.097874534 | 0.097874534 |
| Neuronal acetylcholine receptor; alpha3/beta2 | 0.273748951 | | | | | | | 0.273748951 |
| Neuronal acetylcholine receptor; alpha3/beta4 | 0.559324248 | | | 0.111501865 | | 0.111501865 | 0.097874534 | 0.880202512 |
| Neuronal acetylcholine receptor; alpha4/beta2 | 0.559324248 | | | 0.459066154 | | 0.482634269 | 0.484896363 | 1.985921034 |
| Neuropeptide Y receptor type 1 | | 0.06423878 | | | | | | 0.06423878 |
| Neuropeptide Y receptor type 5 | | | | | 0.100578902 | | | 0.100578902 |
| Nitric oxide synthase, inducible | 0.074564954 | | | | 0.100578902 | | | 0.175143856 |
| Nitric-oxide synthase, brain | 0.074564954 | | | | 0.100578902 | | | 0.175143856 |
| Nitric-oxide synthase, endothelial | 0.074564954 | | | | 0.100578902 | | | 0.175143856 |
| Norepinephrine transporter | | | | | | | 0.097874534 | 0.097874534 |
| Nuclear receptor subfamily 4 group A member 1 | | | | | | | 0.097874534 | 0.097874534 |
| Oligosaccharyl transferase 48 kDa subunit | | | 0.082221517 | | | | | 0.082221517 |
| Orexin receptor 1 | | | | | 0.100578902 | | | 0.100578902 |
| Orexin receptor 2 | | | | | 0.100578902 | | | 0.100578902 |
| Oxytocin receptor | | | | | 0.100578902 | | | 0.100578902 |
| P2X purinoceptor 4 | | | | | | 0 | | 0 |
| P2X purinoceptor 7 | | | | | 0.100578902 | | | 0.100578902 |
| PDZ-binding kinase | | | | 0.111501865 | | 0.111501865 | | 0.22300373 |
| Peptide N-myristoyltransferase 1 | | 0.06423878 | | | | | | 0.06423878 |
| P-glycoprotein 1 | 0.074564954 | 0.217761241 | 0.658856029 | 0.111501865 | | 0.111501865 | 0.097874534 | 1.272060489 |
| PH domain leucine-rich repeat-containing protein phosphatase 2 | | | | | | | 0.097874534 | 0.097874534 |
| Phosphatidylinositol-5-phosphate 4-kinase type-2 gamma | | | | | 0.100578902 | | | 0.100578902 |
| Phosphodiesterase 10A | | | 0.082221517 | | 0.100578902 | | | 0.182800419 |
| Phosphodiesterase 1A | 0.122098887 | | | | | | | 0.122098887 |
| Phosphodiesterase 1C | | | 0.082221517 | | | | | 0.082221517 |
| Phosphodiesterase 4A | | | | | 0.108770969 | | | 0.108770969 |
| Phosphodiesterase 4B | | 0.06423878 | | | 0.100578902 | | | 0.164817682 |
| Phosphodiesterase 4D | 0.074564954 | | 0.082221517 | | 0.100578902 | | | 0.257365373 |
| Phosphodiesterase 5A | | 0.06423878 | | | 0.133337961 | | | 0.197576741 |
| Phosphodiesterase 8B | | | 0.082221517 | | | | | 0.082221517 |
| Phosphodiesterase 9A | | | 0.082221517 | | | | | 0.082221517 |
| PI3-kinase p110-alpha subunit | 0.074564954 | 0.06423878 | 0.082221517 | | 0.100578902 | | | 0.321604153 |
| PI3-kinase p110-delta subunit | | 0.06423878 | | | | | | 0.06423878 |
| Plasma kallikrein | | | | | 0.100578902 | | | 0.100578902 |
| Poly [ADP-ribose] polymerase 2 | | | | 0.111501865 | | 0 | | 0.111501865 |
| Poly [ADP-ribose] polymerase 3 | | | | | 0.100578902 | | | 0.100578902 |
| Poly [ADP-ribose] polymerase-1 | | | | | 0.100578902 | | | 0.100578902 |
| Progesterone receptor | | | | | 0.100578902 | | | 0.100578902 |
| Prolyl endopeptidase | | | | 0.111501865 | 0.100578902 | 0 | | 0.212080767 |
| Prostanoid EP1 receptor | 0.074564954 | | | | | | | 0.074564954 |
| Prostanoid EP3 receptor | | | | | 0.100578902 | | | 0.100578902 |
| Proteasome Macropain subunit MB1 | | | | 0 | | | | 0 |
| Protein arginine N-methyltransferase 6 | | | | | | | 0.097874534 | 0.097874534 |
| Protein arginine N-methyltransferase 8 | | | | | | | 0.097874534 | 0.097874534 |
| Protein farnesyltransferase | | | | 0 | 0.100578902 | | | 0.100578902 |
| Protein kinase C alpha | 0.074564954 | | | 0.111501865 | | 0.111501865 | | 0.297568684 |
| Protein kinase C beta | | 0.06423878 | 0.082221517 | 0.111501865 | | 0.111501865 | | 0.369464027 |
| Protein kinase C delta | 0.074564954 | | | 0.111501865 | | 0.111501865 | | 0.297568684 |
| Protein kinase C epsilon | 0.074564954 | | | 0.111501865 | | 0.111501865 | 0.097874534 | 0.395443218 |
| Protein kinase C eta | 0.074564954 | | | 0.111501865 | | 0.111501865 | | 0.297568684 |
| Protein kinase C gamma (by homology) | | | | | 0.100578902 | | | 0.100578902 |
| Protein kinase C iota | 0.074564954 | | | | | | | 0.074564954 |
| Protein kinase C mu | 0.074564954 | | | | | | | 0.074564954 |
| Protein kinase C theta | 0.074564954 | | | 0.111501865 | | 0.111501865 | | 0.297568684 |
| Protein kinase N1 | | | | | | | 0.097874534 | 0.097874534 |
| Protein kinase N2 | | | | | 0.100578902 | | 0.097874534 | 0.198453436 |
| Protein tyrosine phosphatase receptor type C-associated protein | | | | 0.198389229 | | 0.198389229 | 0.559164811 | 0.955943269 |
| Protein-arginine N-methyltransferase 1 | | | | | | | 0.097874534 | 0.097874534 |
| Proteinase-activated receptor 2 | | | | | 0.100578902 | | | 0.100578902 |

| | | | | | | | | |
|---|---|---|---|---|---|---|---|---|
| Proto-oncogene tyrosine-protein kinase MER | | 0.06423878 | | | | | 0.097874534 | 0.162113314 |
| Proto-oncogene tyrosine-protein kinase ROS | | 0.06423878 | | | | | | 0.06423878 |
| Putative hydrolase RBBP9 | 0.074564954 | | | | | | | 0.074564954 |
| Pyruvate dehydrogenase kinase isoform 1 | 0.074564954 | | | | | | | 0.074564954 |
| Quinone reductase 1 | | | | 0.111501865 | | 0 | | 0.111501865 |
| Quinone reductase 2 | | | | | 0.100578902 | | | 0.100578902 |
| Receptor protein-tyrosine kinase erbB-2 | 0.074564954 | | 0.082221517 | | | | | 0.156786471 |
| Receptor protein-tyrosine kinase erbB-4 | | | 0.082221517 | | | | | 0.082221517 |
| Renin | | 0.06423878 | | | | | | 0.06423878 |
| Rho-associated protein kinase | | | | 0.111501865 | | 0 | 0.097874534 | 0.209376399 |
| Rho-associated protein kinase 1 | | | | 0.111501865 | 0.100578902 | 0.111501865 | 0.097874534 | 0.421457166 |
| Rho-associated protein kinase 2 | 0.074564954 | 0.06423878 | | 0.111501865 | 0.100578902 | 0.111501865 | 0.097874534 | 0.5602609 |
| Ribosomal protein S6 kinase 1 | 0.074564954 | | | | | | | 0.074564954 |
| Ribosomal protein S6 kinase alpha 5 | | | | | | | 0.097874534 | 0.097874534 |
| Serine/threonine-protein kinase 33 | 0.074564954 | | | | | | | 0.074564954 |
| Serine/threonine-protein kinase AKT | 0.074564954 | 0.06423878 | 0.164443033 | 0.111501865 | | 0.111501865 | | 0.526250498 |
| Serine/threonine-protein kinase AKT2 | | | 0.082221517 | | | | | 0.082221517 |
| Serine/threonine-protein kinase Aurora-A | 0.074564954 | 0.06423878 | | | 0.100578902 | | | 0.239382636 |
| Serine/threonine-protein kinase Aurora-B | | 0.06423878 | 0.082221517 | | | | | 0.146460297 |
| Serine/threonine-protein kinase B-raf | 0.074564954 | | | | | | | 0.074564954 |
| Serine/threonine-protein kinase Chk1 | | | 0.082221517 | 0 | | 0.111501865 | | 0.193723382 |
| Serine/threonine-protein kinase D2 | 0.074564954 | | | | | | | 0.074564954 |
| Serine/threonine-protein kinase haspin | | | | 0 | | 0 | 0.097874534 | 0.097874534 |
| Serine/threonine-protein kinase mTOR | 0.074564954 | 0.06423878 | 0.082221517 | | | | | 0.221025251 |
| Serine/threonine-protein kinase NEK2 | 0.074564954 | | | | | | | 0.074564954 |
| Serine/threonine-protein kinase PAK 4 | | | 0.082221517 | | | | | 0.082221517 |
| Serine/threonine-protein kinase PIM1 | 0.074564954 | 0.06423878 | | 0 | 0.100578902 | | 0.097874534 | 0.33725717 |
| Serine/threonine-protein kinase PIM2 | 0.074564954 | | | 0 | 0.100578902 | | 0.097874534 | 0.27301839 |
| Serine/threonine-protein kinase PIM3 | 0.074564954 | | | | | | | 0.074564954 |
| Serine/threonine-protein kinase PLK1 | 0.074564954 | | | | | | | 0.074564954 |
| Serine/threonine-protein kinase PRKX | | | | | | | 0.097874534 | 0.097874534 |
| Serotonin 1a (5-HT1a) receptor | 0.084065592 | | 0.082221517 | 0.387931035 | 0.100578902 | 0.340526084 | 0.592070876 | 1.587394005 |
| Serotonin 1b (5-HT1b) receptor | | | | 0.143102156 | | 0.143102156 | 0.246150751 | 0.532355063 |
| Serotonin 1d (5-HT1d) receptor | | 0.06423878 | | 0.143102156 | | 0.158886034 | 0.246150751 | 0.612377721 |
| Serotonin 1e (5-HT1e) receptor | | | | | | | 0.097874534 | 0.097874534 |
| Serotonin 1f (5-HT1f) receptor | | | | 0.111501865 | | 0.111501865 | 0.097874534 | 0.320878264 |
| Serotonin 2a (5-HT2a) receptor | 0.074564954 | 0.06423878 | | 0.316875924 | | 0.301029825 | 0.534504957 | 1.29121444 |
| Serotonin 2a (5-HT2a) receptor (by homology) | | | 0.082221517 | | 0.100578902 | | | 0.182800419 |
| Serotonin 2b (5-HT2b) receptor | | | 0.082221517 | 0.245758011 | | 0.269486399 | 0.443883112 | 1.041349039 |
| Serotonin 2c (5-HT2c) receptor | | | | 0.15098181 | 0.100578902 | 0.158886034 | 0.246150751 | 0.656597497 |
| Serotonin 3a (5-HT3a) receptor | | 0.06423878 | 0.082221517 | 0.111501865 | | 0.111501865 | 0.097874534 | 0.467338561 |
| Serotonin 4 (5-HT4) receptor | | 0.06423878 | 0.082221517 | 0.111501865 | | 0 | 0.097874534 | 0.355836696 |
| Serotonin 5a (5-HT5a) receptor | | | | 0.166798772 | | 0.174646372 | 0.443883112 | 0.785328256 |
| Serotonin 6 (5-HT6) receptor | | | 0.082221517 | 0.174646372 | 0.100578902 | 0.182601417 | 0.509599763 | 1.04964797 |
| Serotonin 7 (5-HT7) receptor | 0.074564954 | | | 0.356368197 | 0.100578902 | 0.380063355 | 0.6004118 | 1.511987208 |
| Serotonin transporter | 0.074564954 | | | 0.229974302 | | 0.277371341 | 0.312021838 | 0.893932435 |
| Sigma opioid receptor | 0.074564954 | | | 0.111501865 | | 0.111501865 | 0.097874534 | 0.395443218 |
| Sodium channel protein type I alpha subunit | | | 0.082221517 | | | | | 0.082221517 |
| Sodium channel protein type IV alpha subunit | | | | | | | 0.097874534 | 0.097874534 |
| Sodium channel protein type V alpha subunit | | | 0.082221517 | | | | | 0.082221517 |
| Sodium channel protein type X alpha subunit (by homology) | | | | | 0.100578902 | | | 0.100578902 |
| Sodium/glucose cotransporter 1 | | | | | 0.100578902 | | | 0.100578902 |
| Somatostatin receptor 1 | | | | 0.111501865 | | 0.111501865 | | 0.22300373 |
| Somatostatin receptor 4 | | | | 0.111501865 | | 0.111501865 | | 0.22300373 |
| Somatostatin receptor 5 | | 0.06423878 | | | | | | 0.06423878 |
| Squalene synthetase (by homology) | | | | | | | 0.097874534 | 0.097874534 |
| Telomerase reverse transcriptase | 0.074564954 | | | | | | | 0.074564954 |
| TGF-beta receptor type I | 0.074564954 | | | | 0.100578902 | | | 0.175143856 |
| Thrombin | | 0.06423878 | 0.082221517 | | | | | 0.146460297 |
| Thrombin and coagulation factor X | | | | | 0.100578902 | | | 0.100578902 |
| Thromboxane A2 receptor | 0.074564954 | | | | | | 0.097874534 | 0.172439488 |
| Thromboxane-A synthase | | 0.24413516 | | | | | | 0.24413516 |
| Thyrotropin-releasing hormone receptor (by homology) | | | | | 0.100578902 | | | 0.100578902 |
| Transforming growth factor beta-1 | 0.074564954 | | | | | | | 0.074564954 |
| Translocator protein | | | | 0.111501865 | | 0.111501865 | 0.097874534 | 0.320878264 |
| Translocator protein (by homology) | | | | | 0.100578902 | | | 0.100578902 |
| Trypsin I | | 0.06423878 | | | | | | 0.06423878 |
| Trypsin I (by homology) | | | 0.082221517 | | | | | 0.082221517 |
| Trypsin III (by homology) | | | 0.082221517 | | | | | 0.082221517 |
| Tryptase beta-1 | | 0.06423878 | | | | | | 0.06423878 |
| Tryptophan 2,3-dioxygenase | | | | | 0.100578902 | | | 0.100578902 |
| Tyrosine 3-hydroxylase | | | | 0.15098181 | | 0.143102156 | 0.097874534 | 0.3919585 |
| Tyrosine kinase non-receptor protein 2 | | 0.06423878 | 0.082221517 | | | | | 0.146460297 |
| Tyrosine-protein kinase ABL | 0.074564954 | | | | | | | 0.074564954 |
| Tyrosine-protein kinase BTK | | | 0.082221517 | | | | | 0.082221517 |
| Tyrosine-protein kinase CSK | | 0.06423878 | 0.082221517 | | | | | 0.146460297 |
| Tyrosine-protein kinase FYN | | | | | | 0 | | 0 |
| Tyrosine-protein kinase JAK1 | | 0.06423878 | 0.082221517 | 0.111501865 | | 0.111501865 | | 0.369464027 |
| Tyrosine-protein kinase JAK2 | | 0.06423878 | 0.082221517 | 0.111501865 | | 0.111501865 | 0.097874534 | 0.467338561 |
| Tyrosine-protein kinase JAK3 | | 0.06423878 | 0.082221517 | 0.111501865 | | 0.111501865 | | 0.369464027 |
| Tyrosine-protein kinase LCK | | 0.06423878 | 0.082221517 | | | 0 | | 0.146460297 |
| Tyrosine-protein kinase receptor FLT3 | | 0.06423878 | 0.082221517 | | | | 0.097874534 | 0.244334831 |
| Tyrosine-protein kinase receptor TYRO3 | | 0.06423878 | | | | | 0.097874534 | 0.162113314 |

| | | | | | | | | |
|---|---|---|---|---|---|---|---|---|
| Tyrosine-protein kinase receptor UFO | | 0.06423878 | | | | | 0.097874534 | 0.162113314 |
| Tyrosine-protein kinase SRC | 0.074564954 | 0.06423878 | 0.082221517 | 0.111501865 | | 0.111501865 | | 0.444028981 |
| Tyrosine-protein kinase SYK | 0.074564954 | 0.06423878 | 0.082221517 | | 0.100578902 | | | 0.321604153 |
| Tyrosine-protein kinase TYK2 | | 0.06423878 | 0.082221517 | 0.111501865 | | 0.111501865 | | 0.369464027 |
| Tyrosine-protein kinase YES | | 0.06423878 | 0.082221517 | | | | | 0.146460297 |
| Tyrosyl-tRNA synthetase | | | | 0 | | 0.111501865 | | 0.111501865 |
| Urokinase-type plasminogen activator | | | 0.082221517 | | | | | 0.082221517 |
| Urotensin II receptor | 0.074564954 | | 0.082221517 | | | | | 0.156786471 |
| Vascular endothelial growth factor receptor 2 | 0.074564954 | | | | | | | 0.074564954 |
| Vesicular acetylcholine transporter | | | 0.082221517 | | | | | 0.082221517 |
| Voltage-gated potassium channel subunit Kv1.5 | | | | | 0.100578902 | | | 0.100578902 |
| Xaa-Pro aminopeptidase 1 | | | | | 0.100578902 | | | 0.100578902 |
| Xaa-Pro aminopeptidase 2 (by homology) |  |  |  |  | 0.100578902 |  |  | 0.100578902 |
| **Grand Total** | **11.35145403** | **6.804897641** | **9.293199169** | **12.91777542** | **12.63458417** | **12.4244469** | **16.97781013** | **82.40416745** |
