## supplemental data 1a for "Disease Modifying Osteoarthritis Drug Discovery Using A Temporal Phenotypic Reporter In 3D Aggregates of Primary Human Chondrocytes"

| SwissTargetPrediction | Sum of probability |
|---|---|
| Dopamine D2 receptor (by homology) | 2.851672 |
| Dopamine D1 receptor | 2.137331 |
| Neuronal acetylcholine receptor; alpha4/beta2 | 1.985921 |
| Dopamine D3 receptor | 1.749336 |
| Serotonin 1a (5-HT1a) receptor | 1.587394 |
| Serotonin 7 (5-HT7) receptor | 1.511987 |
| Mu opioid receptor | 1.377361 |
| Serotonin 2a (5-HT2a) receptor | 1.291214 |
| P-glycoprotein 1 | 1.27206 |
| Acetylcholinesterase | 1.074565 |
| Serotonin 6 (5-HT6) receptor | 1.049648 |
| Serotonin 2b (5-HT2b) receptor | 1.041349 |
| Dopamine transporter | 1.029377 |
| Adenosine A2b receptor | 1 |
| Protein tyrosine phosphatase receptor type C-associated protein | 0.955943 |
| Serotonin transporter | 0.893932 |
| Neuronal acetylcholine receptor; alpha3/beta4 | 0.880203 |
| Dopamine D4 receptor | 0.865448 |
| Dopamine transporter (by homology) | 0.84436 |
| HERG | 0.830416 |
| Serotonin 5a (5-HT5a) receptor | 0.785328 |
| Alpha-1a adrenergic receptor (by homology) | 0.707984 |
| Multidrug and toxin extrusion protein 1 | 0.705785 |
| Dopamine D5 receptor | 0.693454 |
| Serotonin 2c (5-HT2c) receptor | 0.656597 |
| Alpha-1d adrenergic receptor | 0.639058 |
| Adrenergic receptor alpha-2 | 0.636835 |
| Alpha-2a adrenergic receptor | 0.636835 |
| Alpha-2b adrenergic receptor | 0.636835 |
| Beta-1 adrenergic receptor | 0.630121 |
| Serotonin 1d (5-HT1d) receptor | 0.612378 |
| Butyrylcholinesterase | 0.612333 |
| Alpha-1b adrenergic receptor | 0.590313 |
| Rho-associated protein kinase 2 | 0.560261 |
| Adrenergic receptor beta | 0.541904 |
| Serotonin 1b (5-HT1b) receptor | 0.532355 |
| Serine/threonine-protein kinase AKT | 0.52625 |
| Cyclin-dependent kinase 2/cyclin A | 0.467339 |
| Serotonin 3a (5-HT3a) receptor | 0.467339 |
| Tyrosine-protein kinase JAK2 | 0.467339 |
| Dipeptidyl peptidase IV | 0.459682 |
| Histamine H2 receptor | 0.451933 |
| Tyrosine-protein kinase SRC | 0.444029 |
| Indoleamine 2,3-dioxygenase | 0.421457 |
| Rho-associated protein kinase 1 | 0.421457 |
| Delta opioid receptor | 0.4031 |

| | |
|---|---|
| Coagulation factor VII/tissue factor | 0.395443 |
| Protein kinase C epsilon | 0.395443 |
| Sigma opioid receptor | 0.395443 |
| Tyrosine 3-hydroxylase | 0.391959 |
| Protein kinase C beta | 0.369464 |
| Tyrosine-protein kinase JAK1 | 0.369464 |
| Tyrosine-protein kinase JAK3 | 0.369464 |
| Tyrosine-protein kinase TYK2 | 0.369464 |
| Cyclin-dependent kinase 2 | 0.361807 |
| Inhibitor of nuclear factor kappa B kinase beta subunit | 0.361807 |
| Serotonin 4 (5-HT4) receptor | 0.355837 |
| Serine/threonine-protein kinase PIM1 | 0.337257 |
| Adenosine A1 receptor | 0.321796 |
| ALK tyrosine kinase receptor | 0.321604 |
| PI3-kinase p110-alpha subunit | 0.321604 |
| Tyrosine-protein kinase SYK | 0.321604 |
| Acetylcholine receptor; alpha1/beta1/delta/gamma | 0.320878 |
| Anti-estrogen binding site (AEBS) | 0.320878 |
| Arachidonate 12-lipoxygenase | 0.320878 |
| Arachidonate 15-lipoxygenase | 0.320878 |
| C-C motif chemokine 2 | 0.320878 |
| Dipeptidyl peptidase IX | 0.320878 |
| Fibroblast activation protein alpha | 0.320878 |
| Leukotriene A4 hydrolase | 0.320878 |
| Serotonin 1f (5-HT1f) receptor | 0.320878 |
| Translocator protein | 0.320878 |
| Baculoviral IAP repeat-containing protein 2 | 0.3189 |
| Beta-3 adrenergic receptor | 0.3189 |
| Inhibitor of apoptosis protein 3 | 0.3189 |
| Protein kinase C alpha | 0.297569 |
| Protein kinase C delta | 0.297569 |
| Protein kinase C eta | 0.297569 |
| Protein kinase C theta | 0.297569 |
| Adenosine A2a receptor | 0.281055 |
| Epidermal growth factor receptor erbB1 | 0.27632 |
| Neuronal acetylcholine receptor; alpha2/beta4 | 0.273749 |
| Neuronal acetylcholine receptor; alpha3/beta2 | 0.273749 |
| Serine/threonine-protein kinase PIM2 | 0.273018 |
| Hepatocyte growth factor receptor | 0.257365 |
| MAP kinase p38 alpha | 0.257365 |
| Phosphodiesterase 4D | 0.257365 |
| Tyrosine-protein kinase receptor FLT3 | 0.244335 |
| Thromboxane-A synthase | 0.244135 |
| Serine/threonine-protein kinase Aurora-A | 0.239383 |
| Adenosine A3 receptor | 0.231954 |
| Bromodomain-containing protein 4 | 0.223449 |
| Neuronal acetylcholine receptor protein alpha-7 subunit | 0.223004 |

| | |
|---|---|
| PDZ-binding kinase | 0.223004 |
| Somatostatin receptor 1 | 0.223004 |
| Somatostatin receptor 4 | 0.223004 |
| Alpha-1a adrenergic receptor | 0.221025 |
| Serine/threonine-protein kinase mTOR | 0.221025 |
| Bromodomain-containing protein 2 | 0.215238 |
| Bromodomain-containing protein 3 | 0.215238 |
| Prolyl endopeptidase | 0.212081 |
| Muscarinic acetylcholine receptor M1 (by homology) | 0.209376 |
| Rho-associated protein kinase | 0.209376 |
| Histamine H4 receptor | 0.198453 |
| Melatonin receptor 1B | 0.198453 |
| Muscarinic acetylcholine receptor M2 | 0.198453 |
| Protein kinase N2 | 0.198453 |
| Phosphodiesterase 5A | 0.197577 |
| Interleukin-1 receptor-associated kinase 4 | 0.193723 |
| Serine/threonine-protein kinase Chk1 | 0.193723 |
| Cyclin-dependent kinase 1 | 0.186067 |
| C-C chemokine receptor type 4 | 0.186067 |
| Cathepsin (B and K) | 0.1828 |
| Dopamine D2 receptor | 0.1828 |
| Phosphodiesterase 10A | 0.1828 |
| Serotonin 2a (5-HT2a) receptor (by homology) | 0.1828 |
| Cytochrome P450 2D6 | 0.180096 |
| Cathepsin L | 0.175144 |
| Cyclin-dependent kinase 2/cyclin E1 | 0.175144 |
| Nitric oxide synthase, inducible | 0.175144 |
| Nitric-oxide synthase, brain | 0.175144 |
| Nitric-oxide synthase, endothelial | 0.175144 |
| TGF-beta receptor type I | 0.175144 |
| Monoamine oxidase A | 0.172439 |
| Muscarinic acetylcholine receptor M4 | 0.172439 |
| Thromboxane A2 receptor | 0.172439 |
| Gamma-secretase | 0.164818 |
| Matrix metalloproteinase 1 | 0.164818 |
| Phosphodiesterase 4B | 0.164818 |
| Muscarinic acetylcholine receptor M3 | 0.162113 |
| Proto-oncogene tyrosine-protein kinase MER | 0.162113 |
| Tyrosine-protein kinase receptor TYRO3 | 0.162113 |
| Tyrosine-protein kinase receptor UFO | 0.162113 |
| Macrophage colony stimulating factor receptor | 0.156786 |
| Receptor protein-tyrosine kinase erbB-2 | 0.156786 |
| Urotensin II receptor | 0.156786 |
| Bromodomain testis-specific protein | 0.149733 |
| Inhibitor of NF-kappa-B kinase (IKK) | 0.14913 |
| Bradykinin B1 receptor | 0.14646 |
| Brain adenylate cyclase 1 | 0.14646 |

| | |
|---|---|
| Cathepsin E | 0.14646 |
| C-C chemokine receptor type 3 | 0.14646 |
| C-C chemokine receptor type 5 | 0.14646 |
| DNA-dependent protein kinase | 0.14646 |
| Fibroblast growth factor receptor 1 | 0.14646 |
| Histamine H3 receptor | 0.14646 |
| Histone-lysine N-methyltransferase, H3 lysine-79 specific | 0.14646 |
| Insulin receptor | 0.14646 |
| Mammalian target of Rapamycin (mTORC1) | 0.14646 |
| MAP kinase signal-integrating kinase 2 | 0.14646 |
| Multidrug and toxin extrusion protein 2 | 0.14646 |
| Serine/threonine-protein kinase Aurora-B | 0.14646 |
| Thrombin | 0.14646 |
| Tyrosine kinase non-receptor protein 2 | 0.14646 |
| Tyrosine-protein kinase CSK | 0.14646 |
| Tyrosine-protein kinase LCK | 0.14646 |
| Tyrosine-protein kinase YES | 0.14646 |
| Baculoviral IAP repeat-containing protein 3 | 0.138804 |
| Focal adhesion kinase 1 | 0.138804 |
| Heat shock protein HSP 90-beta | 0.138804 |
| Kinesin-1 heavy chain/ Tyrosine-protein kinase receptor RET | 0.138804 |
| Maternal embryonic leucine zipper kinase | 0.138804 |
| Melanocortin receptor 1 | 0.138804 |
| Melanocortin receptor 3 | 0.138804 |
| Phosphodiesterase 1A | 0.122099 |
| CaM kinase II | 0.111502 |
| Casein kinase I delta | 0.111502 |
| Casein kinase I epsilon | 0.111502 |
| Poly [ADP-ribose] polymerase 2 | 0.111502 |
| Quinone reductase 1 | 0.111502 |
| Tyrosyl-tRNA synthetase | 0.111502 |
| Phosphodiesterase 4A | 0.108771 |
| 11-beta-hydroxysteroid dehydrogenase 1 | 0.100579 |
| Acidic mammalian chitinase | 0.100579 |
| Calpain 1 | 0.100579 |
| Cannabinoid receptor 1 | 0.100579 |
| Carbonic anhydrase II | 0.100579 |
| Cathepsin (V and K) | 0.100579 |
| Cathepsin K | 0.100579 |
| Cathepsin S | 0.100579 |
| CDK8/Cyclin C | 0.100579 |
| Cell division protein kinase 8 | 0.100579 |
| Cyclin-dependent kinase 5/CDK5 activator 1 | 0.100579 |
| Cytochrome P450 11B1 | 0.100579 |
| Cytochrome P450 11B2 | 0.100579 |
| DNA topoisomerase II alpha | 0.100579 |
| Geranylgeranyl transferase type I | 0.100579 |

| | |
|---|---|
| Glucocorticoid receptor | 0.100579 |
| Glycine transporter 1 | 0.100579 |
| Hydroxycarboxylic acid receptor 2 | 0.100579 |
| Insulin-like growth factor I receptor | 0.100579 |
| Legumain | 0.100579 |
| Lysine-specific demethylase 5B | 0.100579 |
| Matrix metalloproteinase 9 | 0.100579 |
| Melatonin receptor 1A | 0.100579 |
| Monoamine oxidase B | 0.100579 |
| Muscarinic acetylcholine receptor M1 | 0.100579 |
| Nerve growth factor receptor Trk-A | 0.100579 |
| Neurokinin 1 receptor | 0.100579 |
| Neuropeptide Y receptor type 5 | 0.100579 |
| Orexin receptor 1 | 0.100579 |
| Orexin receptor 2 | 0.100579 |
| Oxytocin receptor | 0.100579 |
| P2X purinoceptor 7 | 0.100579 |
| Phosphatidylinositol-5-phosphate 4-kinase type-2 gamma | 0.100579 |
| Plasma kallikrein | 0.100579 |
| Poly [ADP-ribose] polymerase 3 | 0.100579 |
| Poly [ADP-ribose] polymerase-1 | 0.100579 |
| Progesterone receptor | 0.100579 |
| Prostanoid EP3 receptor | 0.100579 |
| Protein farnesyltransferase | 0.100579 |
| Protein kinase C gamma (by homology) | 0.100579 |
| Proteinase-activated receptor 2 | 0.100579 |
| Quinone reductase 2 | 0.100579 |
| Sodium channel protein type X alpha subunit (by homology) | 0.100579 |
| Sodium/glucose cotransporter 1 | 0.100579 |
| Thrombin and coagulation factor X | 0.100579 |
| Thyrotropin-releasing hormone receptor (by homology) | 0.100579 |
| Translocator protein (by homology) | 0.100579 |
| Tryptophan 2,3-dioxygenase | 0.100579 |
| Voltage-gated potassium channel subunit Kv1.5 | 0.100579 |
| Xaa-Pro aminopeptidase 1 | 0.100579 |
| Xaa-Pro aminopeptidase 2 (by homology) | 0.100579 |
| Amine oxidase, copper containing | 0.097875 |
| c-Jun N-terminal kinase 1 | 0.097875 |
| Cyclin-dependent kinase 9 | 0.097875 |
| Cyclooxygenase-1 | 0.097875 |
| Cytochrome P450 1A2 | 0.097875 |
| Dipeptidyl peptidase I | 0.097875 |
| Dual-specificity tyrosine-phosphorylation regulated kinase 2 | 0.097875 |
| Hematopoietic prostaglandin D synthase | 0.097875 |
| Kappa Opioid receptor | 0.097875 |
| Leucine-rich repeat serine/threonine-protein kinase 2 | 0.097875 |
| Myosin light chain kinase, smooth muscle | 0.097875 |

| | |
|---|---|
| Neuronal acetylcholine receptor protein alpha-4 subunit | 0.097875 |
| Neuronal acetylcholine receptor; alpha3/alpha6/beta2/beta3 | 0.097875 |
| Norepinephrine transporter | 0.097875 |
| Nuclear receptor subfamily 4 group A member 1 | 0.097875 |
| PH domain leucine-rich repeat-containing protein phosphatase 2 | 0.097875 |
| Protein arginine N-methyltransferase 6 | 0.097875 |
| Protein arginine N-methyltransferase 8 | 0.097875 |
| Protein kinase N1 | 0.097875 |
| Protein-arginine N-methyltransferase 1 | 0.097875 |
| Ribosomal protein S6 kinase alpha 5 | 0.097875 |
| Serine/threonine-protein kinase haspin | 0.097875 |
| Serine/threonine-protein kinase PRKX | 0.097875 |
| Serotonin 1e (5-HT1e) receptor | 0.097875 |
| Sodium channel protein type IV alpha subunit | 0.097875 |
| Squalene synthetase (by homology) | 0.097875 |
| Neurokinin 2 receptor | 0.092752 |
| Cytochrome P450 3A4 | 0.083327 |
| ATP-sensitive inward rectifier potassium channel 1 | 0.082222 |
| Caspase-1 | 0.082222 |
| Ceramide glucosyltransferase | 0.082222 |
| C-X-C chemokine receptor type 7 | 0.082222 |
| dUTP pyrophosphatase | 0.082222 |
| Glutamate NMDA receptor; GRIN1/GRIN2B | 0.082222 |
| Glycogen synthase kinase-3 beta | 0.082222 |
| Gonadotropin-releasing hormone receptor | 0.082222 |
| HLA-DR antigens-associated invariant chain | 0.082222 |
| Macrophage-stimulating protein receptor | 0.082222 |
| MAP kinase p38 beta | 0.082222 |
| Mitogen-activated protein kinase kinase kinase 12 | 0.082222 |
| Mitogen-activated protein kinase kinase kinase 7 | 0.082222 |
| NADPH oxidase 4 | 0.082222 |
| Oligosaccharyl transferase 48 kDa subunit | 0.082222 |
| Phosphodiesterase 1C | 0.082222 |
| Phosphodiesterase 8B | 0.082222 |
| Phosphodiesterase 9A | 0.082222 |
| Receptor protein-tyrosine kinase erbB-4 | 0.082222 |
| Serine/threonine-protein kinase AKT2 | 0.082222 |
| Serine/threonine-protein kinase PAK 4 | 0.082222 |
| Sodium channel protein type I alpha subunit | 0.082222 |
| Sodium channel protein type V alpha subunit | 0.082222 |
| Trypsin I (by homology) | 0.082222 |
| Trypsin III (by homology) | 0.082222 |
| Tyrosine-protein kinase BTK | 0.082222 |
| Urokinase-type plasminogen activator | 0.082222 |
| Vesicular acetylcholine transporter | 0.082222 |
| 3-phosphoinositide dependent protein kinase-1 | 0.074565 |
| CDK9/cyclin T1 | 0.074565 |

| | |
|---|---|
| Complement factor D | 0.074565 |
| C-X-C chemokine receptor type 4 | 0.074565 |
| Histone deacetylase 1 | 0.074565 |
| Histone-lysine N-methyltransferase, H3 lysine-9 specific 3 | 0.074565 |
| Isocitrate dehydrogenase [NADP] cytoplasmic | 0.074565 |
| LIM domain kinase 1 | 0.074565 |
| MAP kinase-activated protein kinase 2 | 0.074565 |
| Methionine aminopeptidase 1 | 0.074565 |
| Prostanoid EP1 receptor | 0.074565 |
| Protein kinase C iota | 0.074565 |
| Protein kinase C mu | 0.074565 |
| Putative hydrolase RBBP9 | 0.074565 |
| Pyruvate dehydrogenase kinase isoform 1 | 0.074565 |
| Ribosomal protein S6 kinase 1 | 0.074565 |
| Serine/threonine-protein kinase 33 | 0.074565 |
| Serine/threonine-protein kinase B-raf | 0.074565 |
| Serine/threonine-protein kinase D2 | 0.074565 |
| Serine/threonine-protein kinase NEK2 | 0.074565 |
| Serine/threonine-protein kinase PIM3 | 0.074565 |
| Serine/threonine-protein kinase PLK1 | 0.074565 |
| Telomerase reverse transcriptase | 0.074565 |
| Transforming growth factor beta-1 | 0.074565 |
| Tyrosine-protein kinase ABL | 0.074565 |
| Vascular endothelial growth factor receptor 2 | 0.074565 |
| Adenosine kinase | 0.064239 |
| Apoptosis regulator Bcl-2 | 0.064239 |
| Apoptosis regulator Bcl-X | 0.064239 |
| Beta secretase 2 | 0.064239 |
| Beta-secretase 1 | 0.064239 |
| Cathepsin D | 0.064239 |
| Cholecystokinin B receptor | 0.064239 |
| C-X-C chemokine receptor type 3 | 0.064239 |
| Cyclin T1 | 0.064239 |
| Cyclin-dependent kinase 4 | 0.064239 |
| Endothelin receptor ET-A | 0.064239 |
| Estrogen receptor alpha | 0.064239 |
| Estrogen receptor beta | 0.064239 |
| Heat shock protein HSP 90-alpha | 0.064239 |
| Histamine H1 receptor | 0.064239 |
| Integrin alpha-IIb/beta-3 | 0.064239 |
| Melanin-concentrating hormone receptor 1 | 0.064239 |
| Melanocortin receptor 4 | 0.064239 |
| Melanocortin receptor 5 | 0.064239 |
| Neuropeptide Y receptor type 1 | 0.064239 |
| Peptide N-myristoyltransferase 1 | 0.064239 |
| PI3-kinase p110-delta subunit | 0.064239 |
| Proto-oncogene tyrosine-protein kinase ROS | 0.064239 |

| | |
|---|---|
| Renin | 0.064239 |
| Somatostatin receptor 5 | 0.064239 |
| Trypsin I | 0.064239 |
| Tryptase beta-1 | 0.064239 |
| Carbonic anhydrase III | 0 |
| Carbonic anhydrase IV | 0 |
| Carbonic anhydrase VA | 0 |
| Carbonic anhydrase VB | 0 |
| Carbonic anhydrase VI | 0 |
| Carbonic anhydrase XII | 0 |
| Carbonic anhydrase XIII | 0 |
| Cyclin-dependent kinase 2/cyclin E | 0 |
| P2X purinoceptor 4 | 0 |
| Proteasome Macropain subunit MB1 | 0 |
| Tyrosine-protein kinase FYN | 0 |
