## supplemental fig1 for "Disease Modifying Osteoarthritis Drug Discovery Using A Temporal Phenotypic Reporter In 3D Aggregates of Primary Human Chondrocytes"

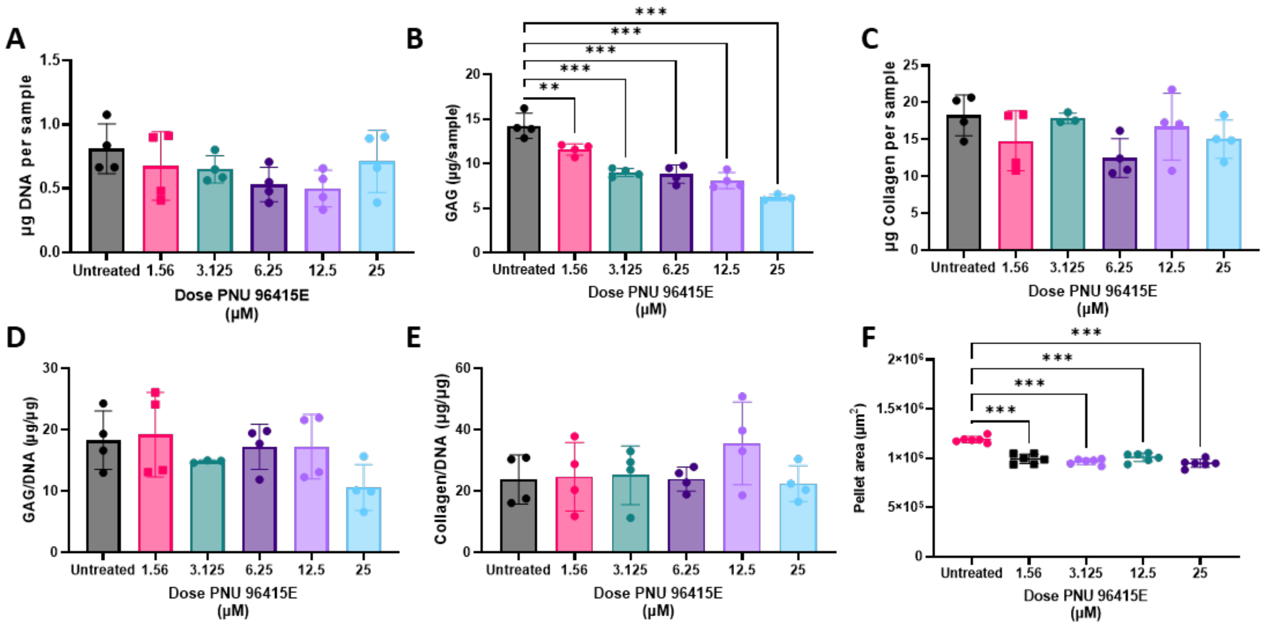


**Supplemental Figure 1. DRD4 antagonist effect on cartilage aggregate biochemistry**

Aggregate DNA (**A**), glycosminoglycan (**B)** and collagen content (**C**) are shown from aggregates treated with the DRD4 antagonist PNU 96415E (0-25 µM). The relative concentrations of GAG/DNA (**D**) and collagen/DNA (**E**) and aggregate size (**F**) are shown. N ≥ 4. Individual replicates or mean of replicates are shown with error bars indicating standard deviation and ** indicating p <0.01, and *** indicating p <0.001 vs. untreated control.
